# High-dimensional mass cytometry reveals stemness state heterogeneity in pancreatic ductal adenocarcinoma

**DOI:** 10.1101/2024.02.27.582358

**Authors:** Egle-Helene Ervin, David Ahern, Feng Liu, Aniko Rendek, Zahir Soonawalla, Udo Oppermann, Philippa Hulley, Siim Pauklin

**Author notes:** **Corresponding author** Siim Pauklin Botnar Research Centre Nuffield Department of Orthopaedics, Rheumatology and Musculoskeletal Sciences University of Oxford Old Road, Headington, Oxford, OX3 7LD United Kingdom.

## Abstract

Stem-like cancer cells harbour high self-renewal capacity, exhibit enhanced tumourigenicity and have been associated with therapy resistance, metastasis and tumour relapse. Therefore, understanding the molecular features of stem-like cells is critical for targeting them effectively and improving treatment outcomes for cancer patients. Several markers have been used to isolate and study the putative stem-like cells of pancreatic ductal adenocarcinoma (PDAC), but the patterns of marker co-expression and overlap between identified individual subpopulations are yet to be comprehensively studied. Here we developed a mass cytometry antibody panel for simultaneous analysis of 33 stemness-associated markers at single-cell resolution. High-dimensional mass cytometry analysis of PDAC cell lines revealed molecularly heterogeneous stemness states and highlighted the role of genotype in determining the cell line-specific stemness signature. Stemness marker expression lie along a continuum in PDAC cell lines and patient samples indicative of stepwise phenotypic transitions. We also identified a subset of PDAC cells co-expressing high levels of Musashi-2, DCLK1 and CXCR4, and harbouring basal-like and EMT transcriptional programmes associated with highly plastic phenotype. This multiplexed analysis uncovers nuance and complexities of the stemness state in the PDAC.

## Introduction

Pancreatic cancer is the seventh leading cause of cancer death worldwide^1^ and has the lowest survival of all common cancers with a 5-year survival rate of 7-12%^2,3^. Due to the increasing incidence and only an incremental improvement in survival, it is projected to become third leading cause of cancer death by 2025 in European countries^1,4^. The most common type of pancreatic cancer is pancreatic ductal adenocarcinoma (PDAC) which accounts for more than 90% of cases, and is associated with poor prognosis due to late diagnosis, aggressive growth and therapy resistance^5,6^. The molecular signature of PDAC involves near ubiquitous activating mutations in *KRAS* (approx. 90% of cases) and inactivating mutations in tumour suppressor genes *CDKN2A* (approx. 90%), *TP53* (50-75%) and *SMAD4* (approx. 55%) which are accompanied by substantial compendium of other, less frequent genomic, epigenomic and transcriptomic alterations^7,8^.

A number of reports have demonstrated that the key features of PDAC underlying the dismal response to treatment and enabling metastatic spread are the phenotypic heterogeneity and plasticity within the tumour cell compartment^9,10^. In particular, the PDAC cell ability to adopt a stem-like state, which is characterised by high self-renewal capacity and enhanced tumourigenicity, has been associated with the formation of metastasis and the tumour relapse^11,12^. Therefore, uncovering the molecular players facilitating the phenotype switching and governing the stem-like state is critical for improving outcomes for the patients with pancreatic cancer. Regarding the latter, numerous markers have been used to isolate and study the stem-like cells of PDAC^13–27^. For example, co-expression of cell surface proteins CD24, CD44 and EpCAM/ESA marks cells with enhanced tumourigenic potential^13^. In addition, cells positive for CD9^14^, CD133^15^, Nestin^24,25^, DCLK1^21^, ALDH^28^ or Musashi-1/2^26^ have also been shown to have stem cell properties. Nevertheless, due to the technical limitations, individual studies of the PDAC stem-like cells have focused only on a handful of factors and there is no overarching concept to explain the identification of many distinct stemness markers^29,30^. Thus, in-depth analysis of the stemness marker expression within the PDAC cell lines and tissue samples is warranted to further understanding of the stem-like state, which in turn enables specific targeting of the stem-like tumour cells and improves patient survival.

To this end, we developed and validated a 41-parameter stemness-centric suspension mass cytometry panel. We used this panel to describe the single-cell marker expression profiles of nine PDAC cell lines, four control cell lines and four PDAC patient tissue samples. The non-adherent (also known as 3D or floating sphere) culture system was used to enrich for stem-like cells and characterise the stemness phenotype. The study revealed considerable heterogeneity between PDAC cell lines and highlighted the role of genotype in determining the cell line-specific stemness signature. Importantly, analysis of the patient samples confirmed that the expression of stemness markers lies along a continuum. Collectively, our results demonstrate that there is no distinct population of universal stem-like cells and support the hypothesis of high plasticity enabling phenotypic transitions which manifest as diverse stemness spectra.

## Results

### Development of stemness-centric mass cytometry panel

Mass cytometry is an advanced molecular biology tool that has a capability to simultaneously quantify over 100 parameters at a single-cell resolution^31^. It has provided key insights into tumour microenvironment of PDAC^32–34^, however, mass cytometry has not yet been used for studying tumour cell compartment. To start to characterise the tumour cell heterogeneity at a protein level and shed light on the stemness state/phenotype, a novel suspension mass cytometry panel (Fig. 1a) was designed with a goal to include all previously identified markers of the PDAC stem-like cells^13–27^ along with the functional markers. The latter serve as readouts of signalling activity, cell cycle phase, epithelial-to-mesenchymal transition (EMT) status and cell viability. In addition, antigens associated with the pluripotency or stem-like cells in other tumour types were incorporated^35–38^.

**Fig. 1:**
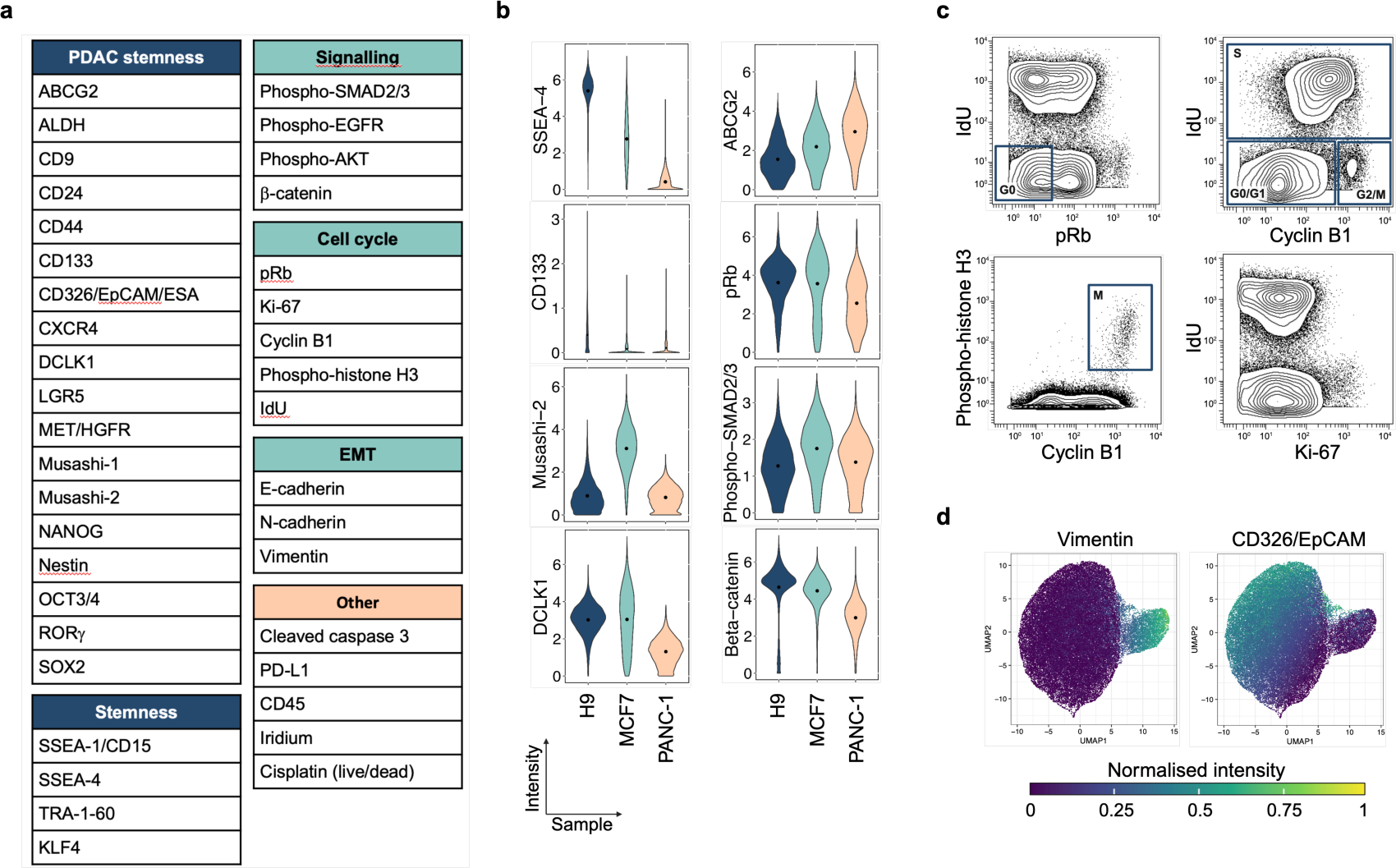
Validation of stemness-centric mass cytometry panel. **a** Stemness-centric suspension mass cytometry panel. For detailed information on all antibodies used in this study, see Supplementary Table 1. **b** Arcsinh-transformed signal intensities of custom-conjugated antibody targets in H9, MCF7 and PANC-1 cell lines. Black lines represent sample mean expression values. **c** Examples of cell cycle marker expression in a sample of PANC-1 cell line. Rectangles represent cell cycle phases. **d** Result of UMAP-based dimensionality reduction on 50,000 cells of H9 cell line. Only EMT markers and CD326/EpCAM were used as input data. Each dot represents a cell and is coloured by the normalised signal intensity of vimentin or CD326/EpCAM. **b-d** Cells were cultured as a monolayer and live, single cells are shown.

Next, metal-conjugated antibodies respective to the above targets were prepared. All custom antibody conjugates were validated using known positive and negative samples, and both custom conjugates and commercially available antibodies were titrated (Fig. 1a, Supplementary Fig. 2 and Supplementary Table 1,2). For example, specific binding of the SSEA-4 and CD133 antibodies was confirmed by using human embryonic stem cell line H9 and pancreatic cancer cell line PANC-1 (Fig. 1b). The former is known to express high levels of SSEA-4^39^ and CD133^40^. In line with the previous studies, cells of the breast cancer cell line MCF7 exhibited the highest expression of Musashi-2 and DCLK1^41–44^. The monoclonal DCLK1 antibody, which recognises mouse and human antigens, was additionally validated on DCLK1-positive mouse embryonic fibroblast cell line NIH3T3 and DCLK1-negative human peripheral blood mononuclear cells (PBMCs)(Supplementary Fig. 2a). The staining pattern of the ABCG2 antibody was concordant with the known features of H9 (ABCG2^low/neg^)^45^ and PANC-1 (ABCG^pos^)^46^ cell lines (Fig. 1b). The cells harbouring high levels of phosphorylated Rb could also be distinguished from the ones where Rb is hypophosphorylated (Fig. 1b,c), and Wnt pathway intracellular signal transducer β-catenin could be detected in, for example, H9, MCF7 and A13A cells but not in unstimulated PBMCs (Fig. 1b and Supplementary Fig. 2a).

In contrast, antibodies detecting RORγ or phosphorylated SMAD2/3, respectively, were excluded from subsequent analysis due to lack of sufficient data to verify their specificity and selectivity (Fig. 1b and Supplementary Fig. 2a). The RORγ is a master transcriptional regulator of CD4 T cell differentiation and a marker of T helper 17 (Th17) cells^47^. However, distinct RORγ-positive Th17 cell population was not observed in the sample of CD4 T cells (Supplementary Fig. 2a). Lastly, the panel allows analysis of cells in different cell cycle phases (Fig. 1c), and discrimination between epithelial (EpCAM^+^, vimentin^-^), mesenchymal (EpCAM^-^, vimentin^+^) and intermediate EMT/MET phenotypes (Fig. 1d). Therefore, this novel, stemness-centric suspension mass cytometry panel has the potential to provide an unprecedented insight into the interplay between stemness, cell cycle, EMT and cell signalling.

### Partly genotype-explained phenotypic differences between pancreatic cancer cell lines

Following successful validation of the antibodies, the panel was employed to characterise the single-cell marker expression profiles of widely used PDAC cell lines (Fig. 2a). Of note, the genotypes of the selected cell lines are representative of the PDAC genomic landscape with the majority of cell lines harbouring an activating mutation in *KRAS* and five cell lines containing an additional alteration in at least one of the PDAC signature genes, namely *TP53*, *CDKN2A* and *SMAD4* (Fig. 2b and Supplementary Table 3)^48^. Considering the aim to describe the stemness phenotype, the culture was enriched for stem-like cancer cells using non-adherent (also known as 3D or floating sphere) culture system^49–52^.

**Fig. 2:**
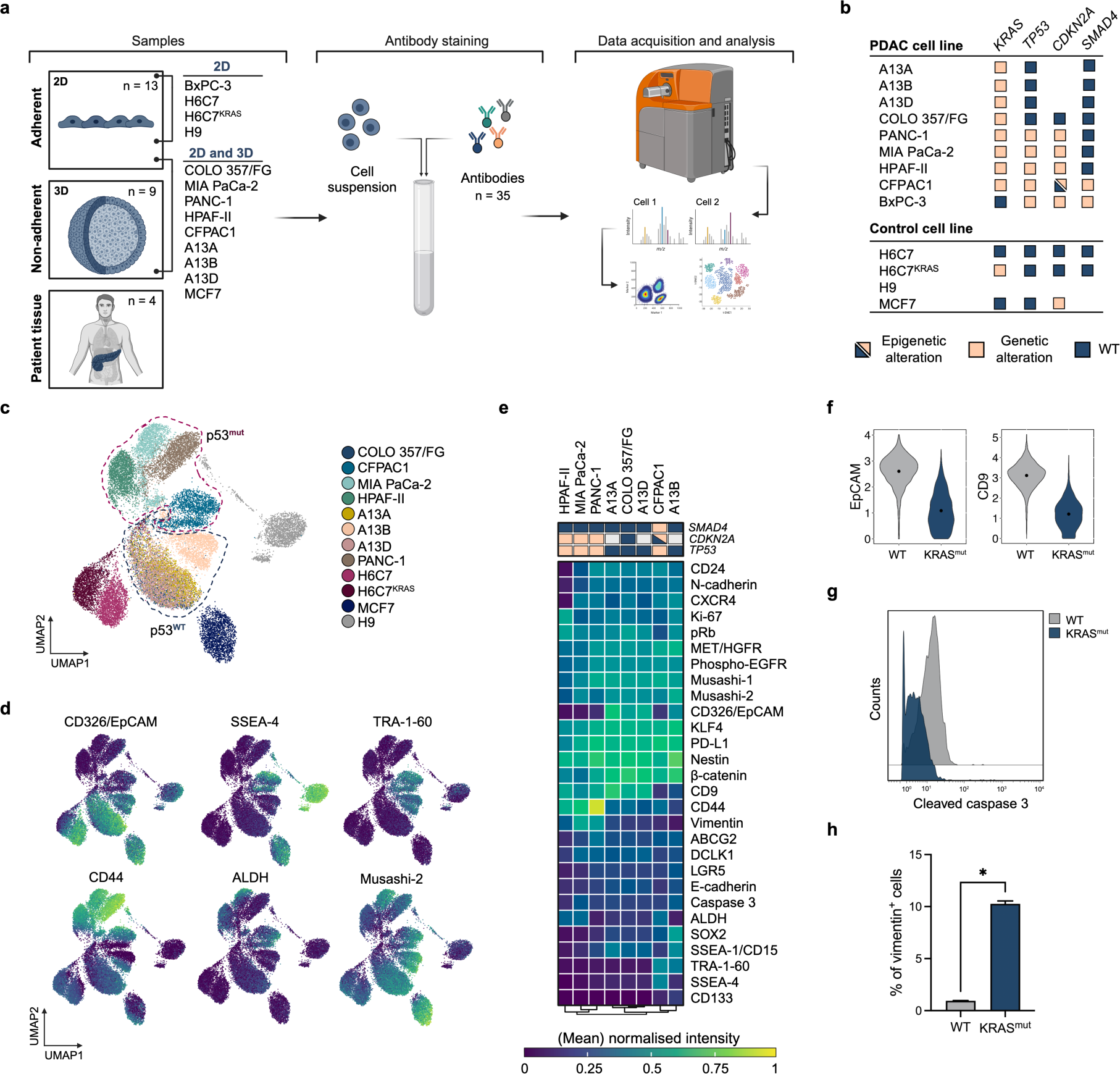
Characterisation of PDAC cell lines cultured in the non-adherent conditions. **a** Schematic of the experimental approach. **b** PDAC cell lines mutational status of *KRAS*, *TP53*, *CDKN2A* and *SMAD4*. Missing values represent genes whose mutational status has not been determined. For detailed information on the genotypes of PDAC cell lines, see Supplementary Table 3. **c** UMAP visualisation of PDAC cell line samples from non-adherent culture. Dimensionality reduction was calculated using subsampled data (2,200 cells/sample) and most variable stemness and EMT markers (see Methods). Each dot represents a cell and is coloured by the cell line. **d** UMAP visualisation as in **c** coloured by normalised expression of indicated markers. **e** Heatmap of mean normalised expression of indicated markers in PDAC cell lines with distinct genetic profiles. **f** Distribution of CD326/EpCAM and CD9 expression in samples of wild-type (WT) or KRAS mutant (KRAS^mut^) H6C7 cell line. Point represents the mean intensity. **g** Histograms of cleaved caspase 3 expression in the samples of wild-type (WT) or KRAS mutant (KRAS^mut^) H6C7 cell line. **h** Percentage of vimentin-positive cells in the samples of wild-type (WT) or KRAS mutant (KRAS^mut^) H6C7 cell line. Data are shown as mean ± SD (n = 2). Two-tailed Student’s t-test, * p-value < 0.05, t-score = 37.32, df = 1. **c-d, f-g** Live, single cells are shown. WT, wild-type.

In total, more than 0.5 × 10^6^ PDAC cells grown in the non-adherent conditions were analysed per biological replicate (Supplementary Table 5). In addition to the PDAC cells, four control cell lines cultured in adherent conditions were studied (Fig. 2a,b). The control cell lines included an immortalised human pancreatic duct epithelial cell line H6C7 (also known as HPDE6c7), H6C7 cell line harbouring oncogenic mutation in *KRAS* (H6C7^KRAS^), human breast cancer cell line MCF7 and human embryonic stem cell line H9. To reveal structure in the data and investigate the role of genetic background in shaping cell phenotype, uniform manifold approximation and projection (UMAP) non-linear dimensionality reduction algorithm was used^53^. The analysis showed that based on the expression of the most variable stemness and EMT markers (Methods), the cells of a single cell line cluster together and separately from the cells of other cell lines (Fig. 2c and Supplementary Fig. 3a).

Therefore, there is significant phenotypic heterogeneity between PDAC cell lines. For instance, HPAF-II, MIA PaCa-2 and PANC-1 cells exhibit lower CD326/EpCAM but higher CD44 levels than the rest of the cells (Fig. 2d,e). While CFPAC1 cells are also CD326/EpCAM^me/lo^, their distinguishing feature is high expression of SSEA-4 and TRA-1-60. Another TRA-1-60^hi^ cell line is A13B which is characterised by enriched expression of SOX2, and like COLO 357/FG, A13A and A13D displays high levels of CD326/EpCAM and medium or low expression of CD24 (Fig. 2d,e and Supplementary Fig. 3b). These phenotypic differences appear to be at least in part associated with the genetic features as the cells carrying a mutant p53 map to a UMAP space distinct from the areas occupied by the cells with a wild-type protein (Fig. 2c). The hierarchical clustering also confirmed that samples harbouring a mutation in *TP53* are more similar to each other than to wild-type (Fig. 2e). Nevertheless, an exception is CFPAC1 which in addition to *TP53* contains a methylated *CDKN2A*, and is the only cell line of the panel with both *KRAS* and *SMAD4* mutations^54^. CFPAC1 groups with A13B due to low CD9 and relatively high TRA-1-60 levels.

Given the clear pattern of mutational profile modulating the cell phenotype, and the previous evidence highlighting the central role of KRAS activation in the PDAC tumour initiation and progression, and thus, in acquisition of a highly tumourigenic state^55^, we investigated whether oncogenic KRAS has an impact on the expression of stemness markers. To this end, H6C7 and an isogenic cell line expressing KRAS^G12V^ (H6C7^KRAS^) were used^56^. On the UMAP plot, H6C7 and H6C7^KRAS^ cells form separate but neighbouring clusters (Fig. 2c). Analysis of the individual markers revealed that H6C7 cells harbouring a mutant KRAS express significantly lower levels of CD326/EpCAM and CD9 (Fig. 2f), and exhibit a reduced number of cells positive for cleaved caspase 3 (Fig. 2g and Supplementary Fig. 4). In contrast, the percentage of cells expressing elevated levels of vimentin is higher in H6C7^KRAS^ sample than in control (Fig. 2h and Supplementary Fig. 4). These results are in line with previous studies reporting the involvement of KRAS in supporting cell survival and facilitating acquisition of a mesenchymal-like phenotype^57,58^. Notwithstanding, the link between the oncogenic activation of KRAS and regulation of the CD9 expression is yet to be established. Of note, six out of eight PDAC cell lines display CD9 levels comparable to the wild-type H6C7 despite activating mutation in *KRAS* (Supplementary Fig. 3b).

In summary, the marker expression profiles of the commonly used PDAC cell lines vary greatly and the mutational background has a profound impact on a cell phenotype.

### Heterogeneous stemness marker expression within pancreatic cancer cell lines

In addition to the differences between cell lines, there is also heterogeneity within individual cell lines. The heterogeneity exists in a form of distinct subpopulations or a low-to-high continuum. As an example of the former, MIA PaCa-2 sample harbours subsets of Nestin^hi^, vimentin^me/hi^ and Nestin^hi^, ALDH^me/hi^ cells (Fig. 3a), respectively, while PANC-1 sample contains a subpopulation of SOX2^hi^, Nestin^hi^ cells. Within A13D and HPAF-II samples, a small group of cells exhibiting elevated levels of SSEA-1 was detected. CFPAC1 cells can be divided into three subgroups based on the expression of CD44. Additionally, the data revealed that the cell line A13B displays greater intrasample diversity than A13A and A13D, and contains a small number of cells with phenotypes that constitute a dominant population in A13A and A13D samples (Supplementary Fig. 3a). Importantly, the A13B, A13A and A13D cell lines are derived from a primary tumour, local metastasis and distant metastasis, respectively, of the same PDAC patient^59,60^. This observation is concordant with the previous report noting greater heterogeneity within A13B and highlighting the selection events occurring during metastatic cascade^60^.

**Fig. 3:**
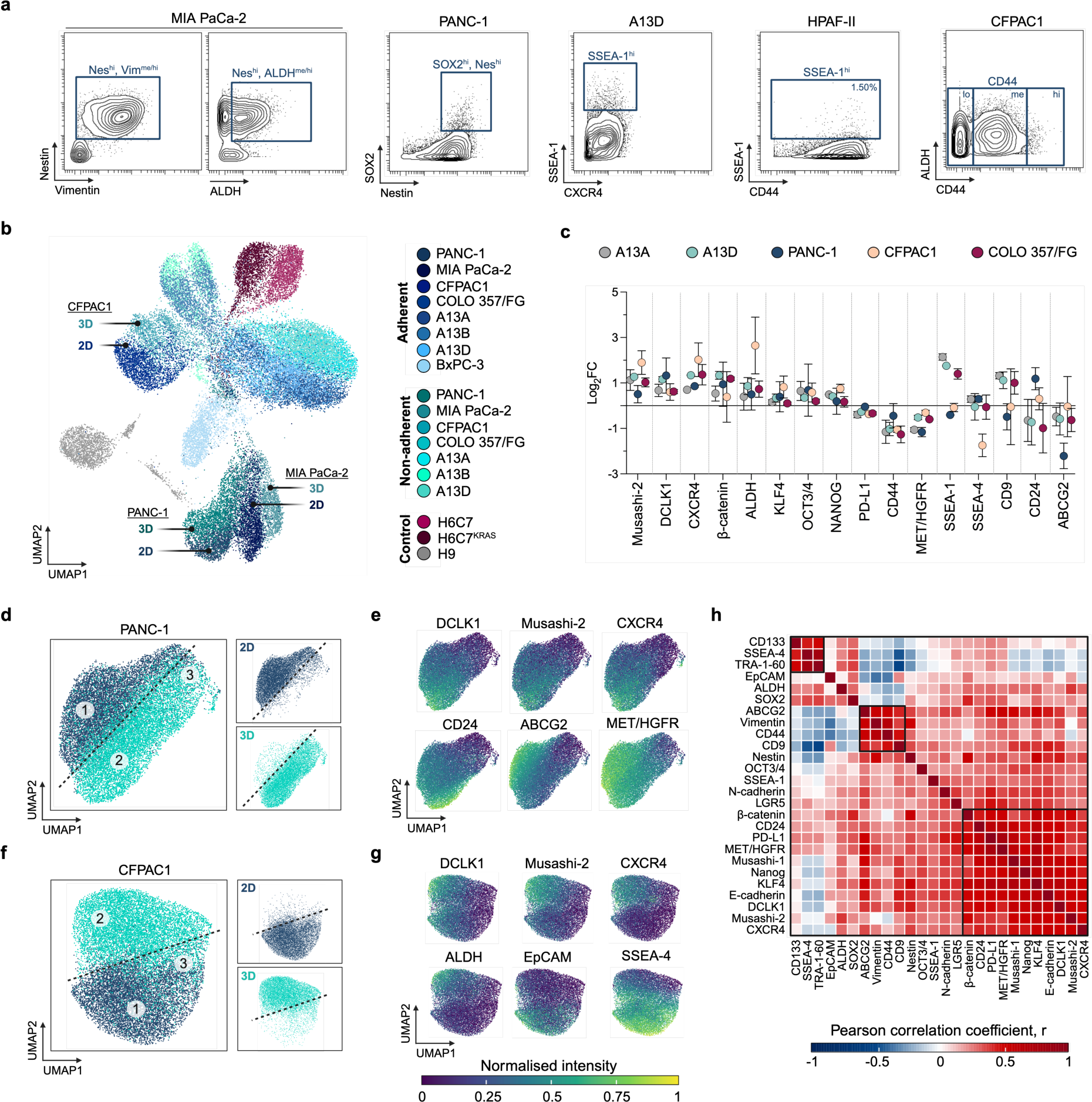
Heterogeneity within and stemness signatures of PDAC cell lines. **a** Two-dimensional contour plots of indicated marker expression in the samples of MIA PaCa-2 (3,850 cells), PANC-1 (10,273 cells), A13D (10,183 cells), HPAF-II (10,215 cells) and CFPAC1 (9,307 cells) cell lines cultured in the non-adherent conditions. Rectangles represent examples of the cell subsets. **b** UMAP visualisation of PDAC and control cell line samples from adherent and non-adherent culture. Dimensionality reduction was calculated using subsampled data (2,200 cells/sample). **c** Relative mean expression levels of indicated markers in non-adherent samples of PDAC cell lines compared to the respective adherent samples. Data are shown as mean ± SD (n = 2). **d** UMAP visualisation of samples of PANC-1 cell line from adherent (2D) and non-adherent (3D) culture. Dimensionality reduction was calculated using subsampled data (9,000 cells/sample). Areas 1-3 exhibit distinct proportions of cells from 2D and 3D culture. **e** UMAP visualisation as in **d** coloured by normalised expression of indicated markers. **f** UMAP visualisation of samples of CFPAC1 cell line from adherent (2D) and non-adherent (3D) culture. Dimensionality reduction was calculated using subsampled data (9,000 cells/sample). Areas 1-3 exhibit distinct proportions of cells from 2D and 3D culture. **g** UMAP visualisation as in **f** coloured by normalised expression of indicated markers. **h** Heatmap of the correlation coefficients for the indicated markers. Correlation analysis was performed on the non-adherent samples of PANC-1, CFPAC1 and COLO 357/FG cell lines (2,220 cells/sample). **a, b, d-g** Live, single cells are shown. **b, d-g** Dimensionality reduction was calculated using most variable stemness and EMT markers (see Methods). Each dot represents a cell and is coloured by the sample (**b, d, f**). FC, fold change.

Notwithstanding the above examples, the expression pattern of the majority of stemness markers forms a spectrum which is exemplified by CD44 expression in HPAF-II and Nestin expression in PANC-1 (Fig. 3a). Likewise, cells with low, intermediate and high levels of Musashi-2, CXCR4, CD24 or CD9 co-exist in individual samples of, for example, COLO-357/FG, PANC-1 and MIA PaCa-2 (Fig. 2d and Supplementary Fig. 3b). These continuous expression values agree with the recent notion that stemness is better described as a dynamic spectrum than as a binary feature^11,61^. Therefore, 1) despite predominantly mapping to a single UMAP area, cells at the opposite ends of a cell line cluster can be significantly dissimilar, and 2) the cells with low and high stemness can be distinguished based on the expression levels of pre-defined markers. The latter is different from a classical stem cell hypothesis stating that the stem cell can be identified solely by the expression (not expression levels) of specific markers that are absent in the non-stem cells^61^.

Regarding the markers characterising the stem-like cells, the fact that there is no single space on the UMAP plot where every cell line is represented suggests that universal, molecularly identical stemness^hi^ phenotype shared by all cell lines does not exist (Supplementary Fig. 3a). Given the above findings, we next sought to describe and compare the stemness signatures of individual PDAC cell lines. Stemness signature is defined as a set of markers that are differentially expressed between stemness^lo^ and stemness^hi^ cells.

### Cell line-specific stemness phenotypes

To delineate the stemness signatures, cells cultured in the non-adherent (3D) system were compared to the respective cells grown in the adherent (2D) conditions (Fig. 2a). The non-adherent culture enriches for stem-like cells, whereas only a small fraction of cells in the adherent conditions are expected to be stem-like^49–52^.

First, UMAP visualisation of all samples demonstrated that neither adherent nor non-adherent samples form a distinct, spatially separated group (Fig. 3b), thereby further supporting the notion that the stemness^hi^ phenotype is cell line-specific. Second, there are phenotypic differences between cells of the same cell line cultured in different (2D versus 3D) conditions as they occupy neighbouring and only minimally overlapping spaces (Fig. 3b). Next, to pinpoint which markers are differentially expressed in the non-adherent sample compared to the respective adherent sample, the mean expression levels were studied. This analysis highlighted Musashi-2, DCLK1 and CXCR4 as markers which are upregulated in all tested non-adherent samples (Fig. 3c). In contrast, CD44 and MET/HGFR exhibit lower expression in non-adherent samples, while effect on the expression of β-catenin, ALDH, SSEA-1, SSEA-4, CD9, CD24 and ABCG2 is cell line-dependent, and levels of KLF4, OCT3/4, NANOG and PD-L1 remain unchanged or change negligibly. Therefore, not all markers previously associated with stemness become upregulated in the non-adherent culture.

We then focused on the individual cell lines at a single-cell resolution to uncover population dynamics and describe the phenotypes enriched in the non-adherent sample. In all three cell lines that were studied in detail, the adherent and non-adherent cells formed a single but clearly polarised cloud where three regions could be identified based on the abundance of cells from individual samples (Fig. 3d, f and Supplementary Fig. 5c). The spatial regions were the following: 1) dominated by the adherent cells, 2) dominated by the non-adherent cells, and 3) containing both adherent and non-adherent cells. This implies that the area 2 is more stem-like than the area 1, since phenotypes mapping to the area 2 are enriched in the non-adherent sample.

In PANC-1 cell line, areas occupied by non-adherent cells are characterised by the high expression of Musashi-2, DCLK1, CXCR4 and CD24, and low or lower levels of ABCG2 and MET/HGFR (Fig. 3e and Supplementary Fig. 6). In CFPAC1 cell line, non-adherent cells exhibit elevated levels of Musashi-1, Musashi-2, DCLK1, CXCR4 and ALDH, but express reduced levels of CD326/EpCAM, SSEA-4 and TRA-1-60 (Fig. 3g and Supplementary Fig. 6). Non-adherent cells of COLO 357/FG cell line demonstrate enhanced expression of Musashi-2 and CXCR4, relatively higher levels of DCLK1, and slightly lower levels of CD44 and ABCG2 than respective adherent cells (Supplementary Fig. 5d and 6). These patterns in the stemness marker expression suggest that while diverse stem-like phenotypes exist, there is a core signature of enhanced levels of Musashi-2, CXCR4 and DCLK1 that is shared between cell lines (Fig. 3c,e,g and Supplementary Fig. 5d). The accessory factors accompanying the core stemness proteins are cell line-specific.

Given the coordinated upregulation of Musashi-2, CXCR4 and DCLK1 in the non-adherent samples, we next asked whether they are co-expressed at a single-cell level. Correlation analysis showed that, among other relationships, there is a moderate positive correlation (Pearson correlation coefficient r of 0.5-0.59) between the expression levels of Musashi-2, CXCR4 and DCLK1 (Fig. 3h). Of note, a strong positive correlation was also observed between the expression of CD44 and vimentin (r = 0.68), and TRA-1-60 and SSEA-4 (r = 0.81). The latter form a co-expression cluster with CD133. In contrast, there is an inverse correlation between CD9 and TRA-1-60 (r = -0.52), and CD9 and SSEA-4 (r = -0.44). Analysis of the individual cell lines confirmed that the markers constituting stemness signature of the respective cell line are co-expressed (Supplementary Fig. 7). For example, Musashi-2, CXCR4, DCLK1 and CD24 group together in PANC-1 sample and there is moderate positive correlation between their expression (0.4 < r ≤ 0.6). In CFPAC1 sample, signal intensities corresponding to Musashi-1, Musashi-2, DCLK1, CXCR4 and ALDH exhibited positive correlation, while weak negative correlation was detected between the expression levels of ALDH and SSEA-4. This suggests that within an individual cell line is a single subset of cells expressing high levels of all stemness signature factors, as opposed to a model whereby individual subpopulations exist each showing enhanced levels of one or two factors.

### Characterisation of stemness marker expression in primary PDAC samples

To characterise the stemness marker expression in primary tumour tissue and investigate whether a subpopulation exhibiting elevated levels of core stemness factors (Musashi-2, CXCR4 and DCLK1) is also present in clinical samples, fresh or frozen surgical tissue from the primary tumours of four pancreatic cancer patients was analysed. In total, 495,815 live single cells were recorded with 38,803-200,685 cells acquired per patient.

First, UMAP dimensionality reduction was run to identify the main cell types and reveal a tumour cell cluster. Based on the canonical marker expression, the epithelial cells (CD326/EpCAM^+^), fibroblasts (vimentin^+^, CD326/EpCAM^-^ and CD45^-^) and immune cells (CD45^+^) were identified within every patient sample with the epithelial cells being the most abundant cell type (Fig. 4a-c and Supplementary Fig. 8a). Second, we focused on the epithelial cells that were grouped into 10 clusters based on the expression of selected stemness, proliferation, EMT, signalling and other (cleaved caspase 3 and PD-L1) markers using unsupervised clustering via Phenograph (Fig. 4d,e and Supplementary Fig. 8b). Cluster 1 contained the highest number of cells and along with cluster 7 was characterised by elevated levels of cleaved caspase 3 but relatively low expression of other antigens. In contrast, clusters 9 and 10 were the smallest and had a distinguishing feature of elevated expression of CD24, CD44 and a set of stemness markers (cluster 9) or high CD24 and E-cadherin levels (cluster 10). Clusters 2 and 8 exhibited enhanced levels of CD9 and NANOG, respectively, while clusters 3 and 4 displayed intermediate expression of MET/HGFR coupled with intermediate levels of CD24 or β-catenin, respectively. Clusters 5 and 6 were among the most proliferative and showed highly similar marker expression profiles of (relatively) high/higher levels of Musashi-1, Musashi-2, β-catenin, DCLK1, Nestin, MET/HGFR and KLF4. The stemness marker expression was slightly lower in the cluster 6 which was also characterised by elevated levels of cleaved caspase 3. Therefore, the stemness marker expression within the epithelial compartment of patient tumours is heterogenous and several subpopulations of varying sizes exhibiting distinct signatures could be detected.

**Fig. 4:**
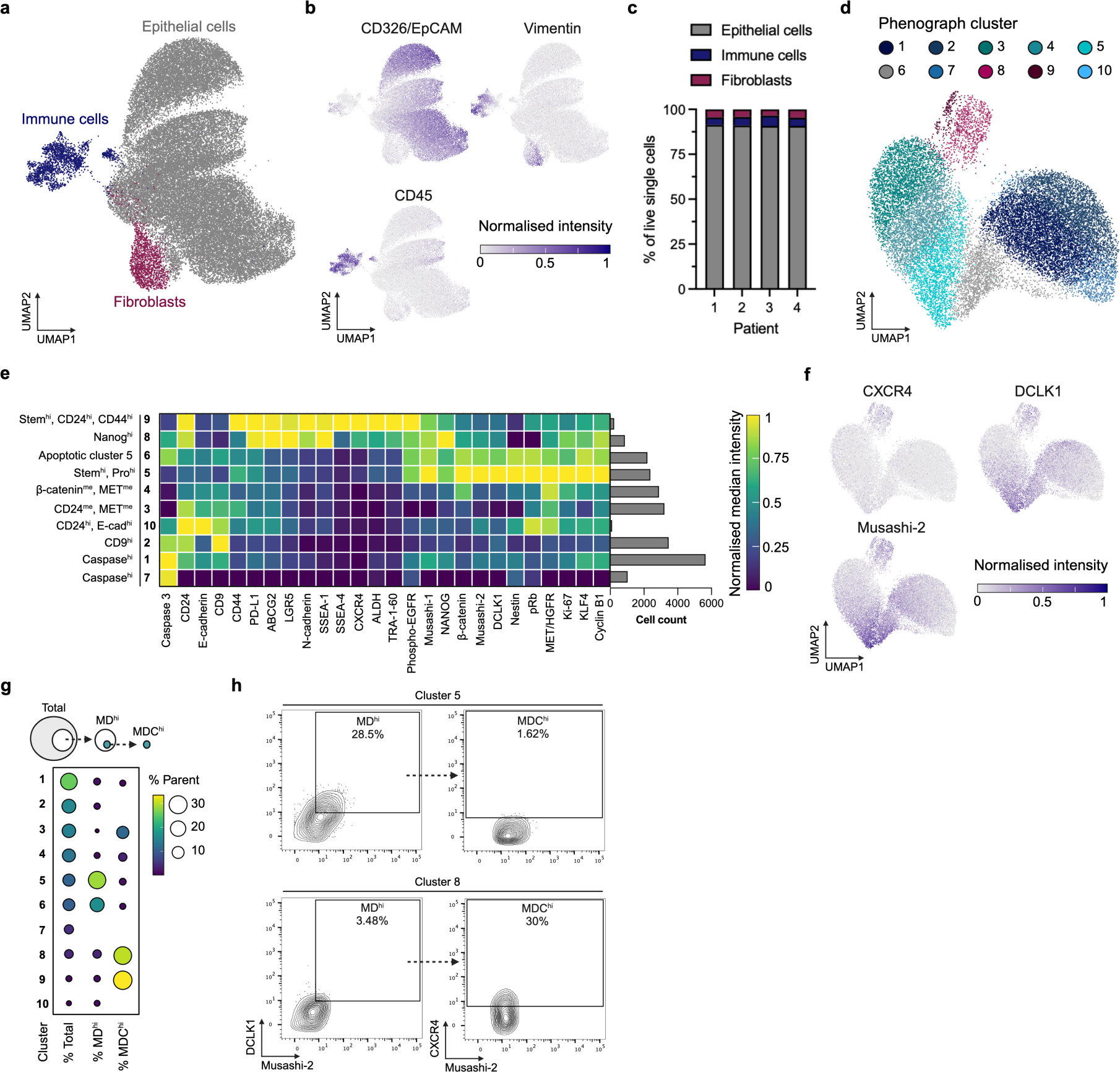
Mass cytometry analysis of primary PDAC samples. **a** UMAP visualisation of all cells from four primary PDAC samples coloured by cell type. Dimensionality reduction was calculated using subsampled data (10,000 cells/patient). **b** UMAP visualisation as in **a** coloured by normalised expression of indicated cell type-specific markers. **c** Frequencies of epithelial cells (CD326/EpCAM^+^), immune cells (CD45^+^) and fibroblasts (vimentin^+^, CD326/EpCAM^-^ and CD45^-^) in individual primary PDAC samples. **d** UMAP visualisation of epithelial cells from four primary PDAC samples coloured by Phenograph cluster. Dimensionality reduction was calculated using subsampled data (5,500 cells/patient). **e** Normalised median expression of indicated markers in the Phenograph clusters of epithelial cells. Cell count represents the number of cells assigned to respective Phenograph cluster. **f** UMAP visualisation as in **d** coloured by normalised expression of indicated markers. **g** Frequencies of MD^hi^ (Musashi-2^hi^ and DCLK1^hi^) and MDC^hi^ (Musashi-2^hi^, DCLK1^hi^ and CXCR4^hi^) cells in Phenograph clusters obtained through sequential gating. Column 1 – proportion of epithelial cells assigned to respective Phenograph cluster, Column 2 – proportion of MD^hi^ cells in respective Phenograph cluster, Column 3 – proportion of MDC^hi^ cells within MD^hi^ population of respective Phenograph cluster. **h** Two-dimensional contour plots illustrating sequential gating of MD^hi^ and MDC^hi^ subpopulations in the Phenograph clusters 5 and 8 based on the indicated marker expression. **a-b, d, f** Dimensionality reduction was calculated using most variable markers (see Methods). Each dot represents a cell. **a-b, d, f, h** Live, single cells are shown.

Third, we aimed to delineate the expression patterns of core stemness factors (Musashi-2, CXCR4 and DCLK1). The unsupervised clustering revealed that the subpopulation exhibiting the highest mean expression of CXCR4 is cluster 9 (stem^hi^, CD24^hi^, CD44^hi^), while the highest Musashi-2 and DCLK1 levels map to cluster 5 (stem^hi^, proliferation^hi^)(Fig. 4e). This was confirmed by UMAP plot where signal intensities corresponding to Musashi-2 and DCLK1 showed significantly similar patterns, and the small number of CXCR4^hi^ cells belonged predominantly to clusters 8 and 9 (Fig. 4f). Importantly, although the cells displaying the highest levels of Musashi-2 and DCLK1 are part of the clusters 5 and 6, a subpopulation of cells in clusters 8 and 9 are also positive for Musashi-2 and DCLK1. To establish whether a triple-high (Musashi-2^hi^, DCLK1^hi^ and CXCR4^hi^) subset exists specifically within clusters 8 and 9, sequential gating was used.

To this end, the gates were set based on cluster 3, a cluster exhibiting the lowest median expression of Musashi-2, DCLK1 and CXCR4 (Fig. 4e), so that less than 1.5% of cells were marker^hi^ (Supplementary Fig. 9a). The analysis corroborated the findings that the highest proportion of double-high (Musashi-2^hi^ and DCLK1^hi^) cells is present in clusters 5 and 6 (Fig. 4g), and clusters 8 and 9 harbour the greatest proportion of triple-high (Musashi-2^hi^, DCLK1^hi^ and CXCR4^hi^) cells. However, the abundance of the latter is low (0.11% of total epithelial cells (Supplementary Fig. 9b)).

In sum, while the triple-high subset could be detected in both established cell lines and primary tumour tissue, the elevated levels of Musashi-2 and DCLK1 tend to be less commonly accompanied by high CXCR4 expression in the clinical samples than in the cell lines. This is indicative of in vitro culture system specifically selecting for triple-high cells or promoting activation of molecular networks driving co-expression of Musashi-2, DCLK1 and CXCR4. Nevertheless, only four clinical samples of early-stage, resectable disease were analysed, thus warranting further studies to capture the whole spectrum of PDAC, and determine which disease stages and subtypes the cell line models represent.

### Transcriptional programmes and clinical relevance of triple-high cells

The enrichment of Musashi-2^hi^, DCLK1^hi^ and CXCR4^hi^ cells in non-adherent culture system suggests these cells harbour molecular programmes critical for stemness properties. To delineate molecular networks underlying their stem-like phenotype, previously published single-cell RNA sequencing (scRNA-seq) data set was used^62^.

The scRNA-seq data set comprised 64,231 epithelial cells (identified based on the canonical marker signature (Supplementary Fig. 10a,b)) from 20 patients. The transcripts of *MSI2*, *DCLK1* and *CXCR4* could be detected (Fig. 5a), and like in the mass cytometry data, cells co-expressing detectable levels of *MSI2*, *DCLK1* and *CXCR4* are a rare subset constituting 0.1% of the epithelial cells. The transcriptomic profile of *MSI2*^+^, *DCLK1*^+^ and *CXCR4*^+^ cells was compared to the rest of cells to reveal differentially expressed genes. In total, 235 significantly upregulated transcripts were identified with the greatest increase in *FN1*, *CXCL14* and *G0S2* mRNA levels (Fig. 5b and Supplementary Fig. 10c).

**Fig. 5:**
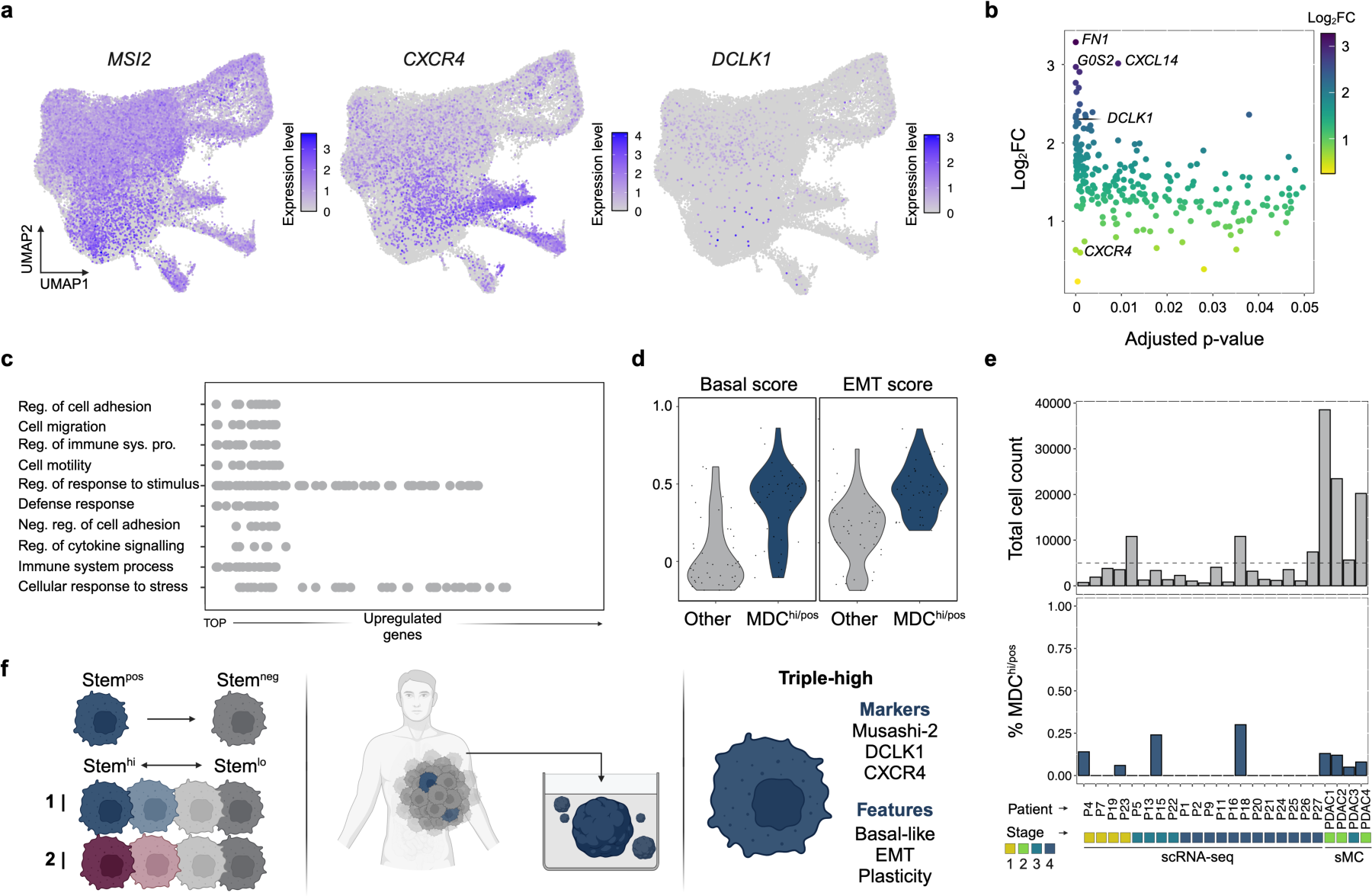
Transcriptional profile of triple-high cells. **a** UMAP visualisation of all epithelial cells from previously published scRNA-seq data set coloured by expression of indicated markers. Dimensionality reduction was calculated using most variable markers (see Methods). **b** Genes significantly (adjusted p-value <0.05) upregulated in triple-high (Musashi-2^hi^, DCLK1^hi^ and CXCR4^hi^) cells compared to the rest of epithelial cells. **c** Functional enrichment analysis of genes in **b**. Top 10 GO biological processes are shown and each dot represents a gene. **d** Basal-like and EMT expression signature scores of MDC^hi^ (Musashi-2^hi^, DCLK1^hi^ and CXCR4^hi^) and other epithelial cells. Subsampled data (43 cells/group) was used and each dot represents a cell. **e** Total cell count and percentage of MDC^hi/pos^ (Musashi-2^hi/pos^, DCLK1^hi/pos^ and CXCR4^hi/pos^) cells in the scRNA-seq and suspension mass cytometry (sMC) samples analysed in the current study. **f** Graphical summary of the following key findings: 1) instead of a universal, molecularly distinct stem-like phenotype, there is heterogeneous stemness continuum; 2) rare primary tumour cellular phenotypes/states may become enriched in in vitro culture, and 3) stemness^hi^ phenotype co-expressing Musashi-2, DCLK1 and CXCR4, and harbouring basal-like and EMT transcriptional programmes was identified. **b-e** Analysis of previously published scRNA-seq data set. FC, fold change; Neg., negative; Pro., processes; Reg., regulation; sMC, suspension mass cytometry; Sys., system

The *FN1* gene encodes fibronectin, a major extracellular matrix (ECM) component, which has recently been implicated in the PDAC tumour progression^63^. The mesenchymal cells enriched in the later disease stages exhibit higher expression of *FN1* and stemness-associated genes^64^. Additionally, an independent study found that the interaction between integrins and tumour-deposited fibronectin is involved in the niche-induced cancer cell stemness^65^. The *CXCL14* gene encodes a member of the CXC chemokine family whose role in the PDAC is relatively unexplored. However, Wente et al. have shown that *CXCL14* mRNA and protein levels are upregulated in the pancreatic cancer compared to the samples of normal pancreas and chronic pancreatitis. CXCL14 is detected at the invasive front of the tumour and its elevated levels increase invasiveness of pancreatic cancer cells^66^. Similarly, the function of *G0S2* gene product in the PDAC is relatively little studied, but it has been demonstrated to be upregulated together with *DCLK1* in the cells that exhibit greater metastatic capacity^67^.

The functional enrichment analysis confirmed that the genes showing elevated expression in *MSI2*^+^, *DCLK1*^+^ and *CXCR4*^+^ cells are associated with terms linked to the cell adhesion, migration and motility (Fig. 5c). Considering the previous reports highlighting the role of mesenchymal and basal-like expression programmes in the pancreatic cancer cell migratory capacity and metastatic propensity^68,69^, the EMT and basal signature scores were calculated for the individual cells. The *MSI2*^+^, *DCLK1*^+^ and *CXCR4*^+^ cells displayed significantly higher expression of EMT and basal signature genes than the rest of the tumour cells (Fig. 5d). Lastly, basal-like PDAC subtype has worse prognosis (Virtual microdissection), but no conclusion on the association between tumour grade and abundance of triple-positive cells could be drawn in the current study due to insufficient sampling (Fig. 5e). To illustrate, 50,000 cells need to be measured to capture 50 cells from a subset constituting 0.1% of the total population.

In conclusion, *MSI2*^+^, *DCLK1*^+^ and *CXCR4*^+^ cells exhibit an increase in the expression of genes implicated in the cell migration and harbour basal transcriptional programme. These molecular features suggest triple-high cells may represent a highly plastic and aggressive subset of PDAC stem-like cells (Fig. 5f)^70^.

## Discussion

Stemness state is an area of great interest in the field of tumour biology due to its involvement in the tumour development, metastatic spread, therapy resistance and relapse^11,71^. Several recent reports have demonstrated that, like the epithelial stem cell heterogeneity and plasticity in the non-cancerous tissues^72^, the population of cancer stem-like cells is heterogeneous and dynamic^73–84^. Here we show that the stemness state heterogeneity and plasticity feature also in the pancreatic cancer as several molecularly non-identical stemness^hi^ states/phenotypes could be detected across PDAC cell lines. The single-cell expression levels of stemness-associated markers populate a continuum in both PDAC cell lines and primary PDAC samples, indicative of stepwise and potentially bidirectional cell state transitions.

With a goal to perform an in-depth analysis of the stemness marker expression within the PDAC cell lines and tissue samples, we opted for suspension mass cytometry which, to our knowledge, has not been applied to study the tumour cell compartment of pancreatic cancer. Suspension mass cytometry has several advantages over the standard single-cell RNA sequencing approaches that are commonly employed for high-content characterisation of cellular identities^85^. For example, it reads out the levels of protein antigens which better reflect the cell functional state than the transcriptomic profile. Additionally, suspension mass cytometry has higher throughput, thereby allowing analysis of rare or low abundant phenotypes^86^. Accordingly, simultaneous analysis of 33 stemness-associated markers at the single-cell resolution provided a glimpse into the functional state heterogeneity of pancreatic cancer stem-like cells and revealed rare co-expression phenotype.

First, the stemness-centric mass cytometry panel along with the adherent and non-adherent culture systems enabled deeper characterisation of widely used cell line models. Our results show that there is significant phenotypic heterogeneity between PDAC cell lines which is linked to cell genetic profile and transcriptomic subtype. To illustrate, HPAF-II, MIA PaCa-2 and PANC-1 cell lines all harbour a mutation in *KRAS*, *TP53* and *CDKN2A*, and exhibit lower CD326/EpCAM but higher CD44 levels than the rest of the cells. MIA PaCa-2 and PANC-1 have been previously shown to represent quasi-mesenchymal transcriptional subtype^87,88^, express EMT programme and display resistance to chemotherapeutic agents^89^. In contrast, a more epithelial cell line COLO 357/FG is lacking a mutation in *TP53* and *CDKN2A*, shows high levels of CD326/EpCAM and relatively low expression of CD24, and is sensitive to chemotherapeutic agents^89^. These findings demonstrate that, expectedly, the molecular differences translate into divergent cell behaviour and highlight the importance of comprehensive characterisation of PDAC cell lines. The latter enables informed and consistent choice of *in vitro* models, thereby helping to unravel the PDAC biology. The single-cell proteomic data presented here complement previous reports describing primarily bulk monolayer cultures^89–92^ and significantly advances our understanding of PDAC cell line characteristics.

Second, while the impact of genotype on the cell phenotype is widely accepted, the role of cancer cell mutational background in modulating its stemness state/phenotype is often overlooked. High-dimensional mass cytometry analysis of cultures enriched for stem-like cells showed that stemness^hi^ state/phenotype is cell line-specific and no universal, molecularly identical stem-like state/phenotype exists. Similarly, the stemness signatures (set of markers that are differentially expressed between stemness^lo^ and stemness^hi^ cells) of the individual cell lines show incomplete overlap. Of note, Gil Vazquez et al. have found similar impact of tumour genotype and subtype on the stem phenotype in colorectal neoplasia^93^. This newly appreciated heterogeneity has a significant impact on our understanding of the stem-like cancer cell biology and their therapeutic targeting. The fact that stem-like features (i.e., self-renewal) are not necessarily associated with a fixed set of previously identified stemness markers implies intricate interplay between cell genetic profile, epigenetic landscape, signalling environment, molecular networks and self-renewal programme in determining the nuanced stem-like state. Thereby, strategies to disrupt the stemness networks have to reflect the complexity and heterogeneity of their target.

Third, although the stem-like cells of individual cell lines are molecularly heterogeneous at the global level, their shared feature is the upregulation of Musashi-2, CXCR4 and DCLK1, which we collectively term core stemness proteins. Musashi-2 is an RNA-binding protein that has been shown to promote pancreatic cancer development, progression and metastasis^41,94–96^. It contributes to drug resistance through a p53-dependent mechanism^97^ and has been used as a reporter to identify the transcriptional programmes maintaining the stem-like state^27^. Musashi-2 expression is associated with poorer differentiation^96^ and it mediates activation of EMT via ZEB1-ERK/MAPK signalling^98^. CXCR4 is a chemokine receptor whose best studied ligand is CXCL12 (also known as stromal derived factor-1, SDF-1)^99^. The CXCL12-CXCR4 axis rescues pancreatic cancer cells from gemcitabine-induced cytotoxicity by stimulating FAK, ERK and AKT signalling pathways, and promoting transcriptional activity of β-catenin and NF-κB^100^. CXCR4 expression is associated with PDAC progression^101^ and malignant features (e.g., desmoplastic reaction and migration)^102,103^. Importantly, CXCR4 has been shown to mark a subpopulation of stem-like pancreatic cancer cells exhibiting enhanced invasive and metastatic properties^15^. DCLK1 is a bifunctional protein with kinase and microtubule-associated protein activity that identifies a rare population of long-lived, quiescent pancreatic progenitors facilitating regeneration and underlying tumour initiation^104,105^. DCLK1-positive cells detected in the preinvasive lesions and pancreatic cancer display features of gastrointestinal tuft cells, express genes linked to the cancer stem cells and harbour an active EMT programme^21,43,106,107^. Accordingly, DCLK1 is a putative marker of pancreatic cancer stem-like cells that is essential for their invasive and metastatic property^108^. It is also highly expressed by circulating tumour cells and metastatic tumours, and its inhibition reverses EMT programme and restores T cell activity^43,109,110^.

Given the previously described functions of the individual core stemness proteins, it is unsurprising that the cells co-expressing Musashi-2, CXCR4 and DCLK1 (triple-high/positive phenotype) are transcriptionally characterised by elevated EMT and basal scores. EMT is a dynamic and reversible process that has been associated with metastasis, tumour-initiating potential, stemness and therapy resistance^111^. Basal cells have recently been demonstrated to represent a highly plastic state with the capacity to facilitate state transitions and promote intratumoural heterogeneity in PDAC^112^. These findings are in accordance with the notion that the stemness, EMT, plasticity, metastasis and therapy resistance are all closely interlinked^111,113^. The identified triple-high/positive stem-like cells may be a highly plastic tumour cell phenotype implicated in the metastatic spread and adaptive therapy resistance of the PDAC. Our data support the plasticity and state transitions surrounding the stem-like cells which are expected to result in the continuum of marker expression and spectrum of intermediate cell states^112,114^.

Future research will help to address the limitations of the current study and answer the outstanding questions. For example, (spatial) analysis of a larger number of patient samples is critical for confirming stemness state heterogeneity, understanding the clinical significance of the triple-high cells and shedding light on the stem-like cell niche/cellular neighbourhood. Mechanistic studies elucidating the origin, drivers, dynamics and functional differences of the heterogeneous stem-like cell subpopulations, and the role of triple-high cells in the tumour development, progression, metastasis and therapy resistance are also crucial next steps.

In conclusion, high-dimensional mass cytometry analysis reveals stemness state heterogeneity and plasticity in the PDAC. Our results add to the accumulating evidence describing stemness as a dynamic spectrum, not as a fixed binary feature. This paradigm shift highlights the need to develop new therapies that simultaneously target the individual cancer cell states and their plasticity machinery.

## Methods

### Cell lines

A13A, A13B and A13D cell lines were kindly provided by Prof. Christine Iacobuzio-Donahue (Memorial Sloan Kettering Cancer Center). COLO 357/FG cell line was obtained from MD Anderson Cancer Center Characterized Cell Line Core Facility. PANC-1, HPAF-II, CFPAC1 and MCF7 cell lines were kindly provided by Prof. Catherine Hogan (Cardiff University). MIA PaCa-2 and BxPC-3 cell lines were kindly provided by Prof. Eric O’Neill (University of Oxford). H6C7 cell line was obtained from Kerafast and H6C7^KRAS^ cell line was kindly provided by Prof. Ming Tsao (University of Toronto). NIH3T3 cell line, PBMCs and CD4 T cells were kindly provided by Prof. Udo Oppermann, Prof. Claudia Monaco and Dr Liye Chen (all University of Oxford), respectively. H9 (WA09) cell line was obtained from WiCell. All cell lines were regularly tested for mycoplasma infection.

### Adherent cell culture

PDAC cell lines A13A, A13B, A13D, COLO 357/FG, PANC-1, HPAF-II, CFPAC1, breast cancer cell line MCF7 and mouse fibroblast cell line NIH3T3 were cultured in Dulbecco’s Modified Eagle Medium (DMEM) supplemented with 10% FBS, 1% Penicillin-Streptomycin, 1% MEM non-essential amino acids solution and 1% MEM vitamin solution (all Thermo Fisher). PDAC cell line MIA PaCa-2 was cultured in DMEM supplemented with 10% fetal bovine serum (FBS), 1% Penicillin-Streptomycin, 1% MEM non-essential amino acids solution, 1% MEM vitamin solution and 2.5% horse serum (all Thermo Fisher). PDAC cell line BxPC-3 was cultured in Roswell Park Memorial Institute (RPMI) 1640 Medium supplemented with 10% FBS (both Thermo Fisher). Human pancreatic duct epithelial cell lines H6C7 and H6C7^KRAS^ cell lines were cultured in keratinocyte serum-free medium (SFM) supplemented with 5 ng/ml human recombinant epidermal growth factor (EGF) and 50 µg/ml bovine pituitary extract (BPE)(all Thermo Fisher). Above cells were split every 4-5 days using 5-10-minute incubation with 0.05% trypsin-EDTA (Thermo Fisher) at 37 °C. Human embryonic stem cell line H9 was cultured on the 6-well plate coated with the Vitronectin (5 μg/ml) and in the Essential 8 Flex medium (both Thermo Fisher). Cells were split every 7 days using 5-minute incubation with 0.5 mM EDTA-PBS (Thermo Fisher) at 37 °C. All cells were cultured in humidified incubator at 37 °C with 5% CO_2_.

### Non-adherent cell culture

PDAC cell lines A13A, A13B, A13D, COLO 357/FG, PANC-1, MIA PaCa-2, HPAF-II, CFPAC1 and breast cancer cell line MCF7 were cultured in non-adherent system of ultra-low attachment vessel and serum-free DMEM/Nutrient Mixture F-12 medium (Sigma-Aldrich) supplemented with 2% B-27, 1% L-glutamine, 1% Penicillin-Streptomycin (all Thermo Fisher) and bFGF (20 ng/ml, Peprotech). Spheroids were split every 5 days using 5-min incubation with TrypLE™ Express (Thermo Fisher) at 37 ℃. Cells were passed through a cell strainer (40 µm, pluriSelect) followed by counting and reseeding single cells at a density of 2,000 cells/ml. Cells were cultured in humidified incubator at 37 °C with 5% CO_2_.

### Patient tissue samples

Primary PDAC tissue samples from a total of four patients were obtained from the Oxford Radcliffe Biobank. Written informed consent was obtained from all participants and study was conducted in compliance with the ethics permission (OCHRe ref: 21/A126, REC reference: 19/SC/0173; OCHRe ref: 19/A176, REC reference: 19/SC/0173). Tissue punch biopsies (size: 5 × 5 mm^2^) were cut immediately after resection and PDAC diagnosis in all samples was confirmed by certified pathologist. Fresh surgical tissue was stored in ice-cold DMEM (Thermo Fisher) before it was minced using scalpel, and cryopreserved in FBS supplemented with 10% dimethyl sulfoxide (DMSO) at -80 °C (samples for genotyping and mass cytometry) or processed directly using Tumor Dissociation Kit (human, Miltenyi Biotec) as described below (fresh sample for mass cytometry). To determine the genetic variants of key PDAC-associated genes (*KRAS*, *TP53*, *SMAD4* and *CDKN2A*), genomic DNA was extracted from cryopreserved patient tissue samples using GenElute Mammalian Genomic DNA Miniprep Kit (Sigma-Aldrich) according to manufacturer’s instructions. Purified genomic DNA (10 ng, eluted in nuclease-free water) was prepared for next generation sequencing using Ion AmpliSeq Library Kit Plus, Ion AmpliSeq Cancer Hotspot Panel v2 and Ion Xpress Barcode Adapters according to manufacturer’s instructions. Sequencing libraries were prepared with Ion 510 & Ion 520 & Ion 530 Kit – Chef using Ion Chef System, sequenced on Ion GeneStudio S5 System (all Thermo Fisher) and aligned to GRCh37 following manufacturer’s recommended data processing pipeline. Customised workflow in Ion Reporter software 5.12 was used for variant calling and annotation.

### Mass cytometry panel

Mass cytometry panel allowed to measure 47 parameters/channels: 38 antigens (antibody targets), 6 barcodes, 1 S phase marker (IdU), 1 intercalator (Iridium) and 1 live/dead marker (Cisplatin)(Supplementary Table 1). Heavy metal-labelled antibodies were purchased from the Standard BioTools or prepared using Maxpar Antibody Labelling Kit (Standard BioTools) according to manufacturer’s instructions. Validation of in-house-labelled antibodies was performed on known marker-negative and marker-positive cells (Supplementary Table 2). Optimal staining concentration was determined using 8-point (2-fold dilution series from 0 to 8 µg/ml) or 3-point (1:200, 1:100 and 1:67) titration of in-house-labelled and commercially available antibodies, respectively. If the 1:200 dilution resulted in a too high signal intensity, lower concentrations were tested. The list of antibodies, clone information, metal isotype tags used and staining concentrations are provided in Supplementary Table 1.

### Processing of cell line samples

Spent culture medium was removed and cells were incubated in a serum-free medium containing Cell-ID™ 127 IdU (25 µM, Standard BioTools) for 30 min at 37 °C. Following a wash with PBS (Thermo Fisher), single-cell suspension was created by incubation in TrypLE™ Express (Thermo Fisher) supplemented with Benzonase nuclease (1:10,000, Santa Cruz Biotechnology) for 10 min at 37 °C. Single-cell suspension was washed with PBS (Thermo Fisher) and incubated in a serum-free medium containing viability reagent Cell-ID™ Cisplatin (1 µM, Standard BioTools) for 10 min at 37 °C. Cisplatin was quenched by washing the cells twice with serum-containing medium (DMEM supplemented with 10% FBS, adherent samples) or Maxpar Cell Staining Buffer (Standard BioTools, non-adherent samples). Cells were fixed in a 4% formaldehyde, methanol-free (Thermo Fisher) for 10 min at room temperature. Fixation solution was removed, cells were resuspended in cold Maxpar Cell Staining Buffer and counted. For mass cytometry staining, using 0.5 or 0.75 × 10^6^ cells per sample was preferred but lower number of cells was used if enough cells could not be obtained.

### Processing of patient tissue samples

Cryopreserved patient tissue samples were thawed rapidly into DMEM. Thawed and fresh samples were dissociated using the Tumor Dissociation Kit (human, Miltenyi Biotec) and tough dissociation protocol on gentleMACS Dissociator (Miltenyi Biotec) according to manufacturer’s instructions. Following dissociation, samples were resuspended in DMEM (Thermo Fisher) and treated with Benzonase nuclease (1:10,000, Santa Cruz Biotechnology) for 5 min at room temperature. Benzonase-containing medium was removed by centrifugation and cells were incubated in a serum-free medium with Cell-ID™ 127 IdU (25 µM, Standard BioTools) for 20 min at 37 °C. Subsequently, viability reagent Cell-ID™ Cisplatin (1 µM, Standard BioTools) was added and cells were incubated for a further 10 min at 37 °C. Cisplatin was quenched by washing the cells once with serum-containing medium and once with Maxpar Cell Staining Buffer. Cells were fixed in a 4% formaldehyde, methanol-free (Thermo Fisher) for 10 min at room temperature. Fixation solution was removed, cells were resuspended in cold Maxpar Cell Staining Buffer and counted. For mass cytometry staining, 0.1-0.5 × 10^6^ cells per sample were used.

### Mass cytometry staining

Cells were prepared as described above, washed with Maxpar Cell Staining Buffer, and stained with CXCR4 antibody for 30 min at 4 °C. Upon incubation, cells were washed with Maxpar Cell Staining Buffer and barcoded using Cell-ID™ 20-Plex Pd Barcoding Kit (Standard BioTools) according to manufacturer’s instructions. Barcoded cells were combined into a composite sample before incubation in Human TruStain FcX Fc Receptor Blocking Solution (BioLegend) for 10 min at room temperature. Blocking solution was removed by centrifugation, volume of cell suspension was adjusted to achieve a concentration of 3 × 10^6^ cells in 100 µl and cells were stained with surface antibody cocktail for 30 min at 4 °C. After the incubation, cells were washed with Maxpar Cell Staining Buffer and chilled on ice for 10 min. For methanol fixation/permeabilisation, ice-cold 90% methanol (Thermo Fisher) was slowly added to cells and cells were incubated in methanol for 30 min on ice. Next, cells were washed twice with Maxpar Cell Staining Buffer supplemented with Benzonase nuclease (1:10,000, Santa Cruz Biotechnology). Volume of cell suspension was adjusted to achieve a concentration of 3 × 10^6^ cells in 100 µl and cells were stained with intracellular antibody cocktail for 30 min at room temperature. Following two washes with Maxpar Cell Staining Buffer, samples were incubated in intercalation solution (1.6% formaldehyde and 62.5 nM Cell-ID Intercalator-Ir (Standard BioTools) in Maxpar PBS (Standard BioTools)) overnight at 4 °C. On the day of acquisition, cells were washed once with Maxpar Cell Staining Buffer and once with Maxpar Water (Standard BioTools), filtered through cell strainer (35 μm, Falcon), resuspended in Maxpar Cell Acquisition Solution with 0.1X EQ beads and acquired on Helios mass cytometer (all Standard BioTools).

### Mass cytometry data analysis

Raw mass cytometry data were bead normalised and debarcoded on CyTOF Software v7.0. Population of live, single cells for further analysis was identified by manual gating on Cytobank as described in Supplementary Fig. 1. A summary of samples and corresponding number of events for all experiments are reported in Supplementary Table 5. Data were subsequently imported into R^115^ environment and batch corrected by scaling to the 95^th^ percentile of the anchor sample (adherent A13A) using BatchAdjust package^116^. Harmonised data were arcsinh transformed (cofactor 5) and downstream analysis was performed using tidytof package^117^. Uniform manifold approximation and projection (UMAP) dimensionality reduction was calculated on downsampled data with the following parameters: set.seed(0), neighbors = 15 and min_dist = 0.5. The following channels, which showed the greatest variation between samples (visual inspection), were used for UMAP analysis: 1) Cell line samples: EpCAM, N-cadherin, SSEA-1, SSEA-4, ALDH, TRA-1-60, CD133, SOX2, Nestin, vimentin, Musashi-1, CXCR4, E-cadherin, DCLK1, LGR5, OCT3/4, CD24, Nanog, ABCG2, CD44, CD9, Musashi-2 and β-catenin; 2) Patient tissue samples (all cells): Ki-67, pRb, cyclin B1, caspase 3 (cleaved), ABCG2, β-catenin, CD9, CD24, CD45, CXCR4, DCLK1, Musashi-1, MET/HGFR, Nestin, EpCAM, E-cadherin, vimentin and PD-L1; 3) Patient tissue samples (epithelial cells): caspase 3 (cleaved), ABCG2, ALDH, β-catenin, CD9, CD24, CD44, CD133, CXCR4, DCLK1, KLF4, LGR5, Musashi-1, Musashi-2, MET/HGFR, Nestin, OCT3/4, SOX2, SSEA-1, SSEA-4, TRA-1-60, E-cadherin, N-cadherin, vimentin and PD-L1. Importantly, since distinct CD44^+^ and CD44^-^ subpopulations within adherent and non-adherent samples of CFPAC1 cell line hindered intersample comparisons, CD44 was excluded from UMAP analysis shown in Fig. 3f,g. Epithelial cells from patient tissue samples were clustered based on the expression of above markers (UMAP channels set number 3) using PhenoGraph algorithm with the following parameters: set.seed(0), num_neighbors = 50 and distance_function = “cosine”. Pearson correlation coefficients were calculated using the R stats package^115^.

### Single-cell RNA-seq data analysis

Single-cell RNA-seq data on 20 treatment-naïve patients were obtained from a study reported by Werba et al. (GEO accession number GSE205013)^118^. Paired reads were aligned to the hg38 reference genome and gene expression matrices were generated using 10x Genomics Cell Ranger 7.0.0. Subsequent analysis on data matrices was performed using Seurat package (v4)^119^. Low quality cells were removed based on the following metrics: genes detected per cell (<400) and UMI counts per cell (<600). Cells with high mitochondrial counts were removed at the clustering step. Raw count data were normalised and variable genes for each sample were identified using NormalizeData and FindVariableFeatures functions, respectively. FastMNN algorithm was used for integration of individual samples. For cell type identification, dimensionality reduction was performed with RunUMAP on highly variable features followed by the clustering of cells based on their global transcriptional profiles using FindNeigbors and FindClusters functions. Cell types were annotated based on the marker genes in Supplementary Fig. 10b. For subsetting triple-positive cells, epithelial cells with feature count higher than 0 for *MSI2*, *DCLK1* and *CXCR4* were considered. Differential expression analysis was performed with FindMarkers function and g:GOSt tool on g:Profiler^120^ web server was used for functional enrichment analysis of differentially expressed genes. Gene expression signature scores were obtained by dividing the sum of normalised and scaled counts of each feature/gene with the number of features/genes constituting the signature. Top 100 basal-like and EMT signature genes defined by Chang-Seng-Yue et al.^121^ and Puram et al.^122^, respectively, were used.

### Visualisation

Plots were created using the GraphPad Prism version 10.0.2, ggplot2^123^ and pheatmap^124^(both R packages). Schematic representations were created and figures were prepared with BioRender.

### Statistics

All statistical tests used in this study are described in detail in the corresponding figure legends. The experiment comparing single-cell profiles of PDAC cell lines grown in adherent versus non-adherent culture was repeated twice (n = 2).

## Data availability

The mass cytometry data are available on Zenodo. The existing, publicly available scRNA-seq data used in this paper are available on GEO with accession number GSE205013.

## Code availability

No custom code was used for processing the data presented in the manuscript.

## Acknowledgements

The authors thank Christine Iacobuzio-Donahue, Catherine Hogan, Eric O’Neill, Ming Tsao, Claudia Monaco and Liye Chen for providing the cell lines, and Jonathan Williams and Sarah Pericli for their advice and assistance with clinical sample genotyping. The work in the laboratory of S.P. was supported by a Cancer Research UK Career Development Fellowship (Grant ID: C59392/A25064) and Pancreatic Cancer UK (Grant/Award Number: 2018RIF_03). E-H.E. was supported by the Kristjan Jaak Scholarship Programme. U.O. was supported by Innovate UK, the National Institute for Health Research Oxford Biomedical Research Centre, Cancer Research UK, the Leducq Epigenetics of Atherosclerosis Network (LEAN) program grant from the Leducq Foundation and the Myeloma Single Cell Consortium.

## Author contributions

E-H.E., D.A., and S.P. designed the project and performed the experiments. E-H.E. and F.L. analysed the data and visualised the results. A.R. and Z.S. provided pathology support and clinical tissue samples. S.P. provided resources. S.P., P.H. and U.O. supervised the project and provided intellectual input. The original draft was written by E-H.E. and S.P. All authors reviewed the manuscript.

## Materials and correspondence

Correspondence and material requests to Siim Pauklin.

