## Supplementary Material for "High-dimensional mass cytometry reveals stemness state heterogeneity in pancreatic ductal adenocarcinoma"

### Contents

#### Supplementary Figures

Supplementary Fig. 1: Gating strategy for mass cytometry data clean-up.
Supplementary Fig. 2: Mass cytometry antibody validation and titration.
Supplementary Fig. 3: Characterisation of PDAC cell lines cultured in the non-adherent conditions.
Supplementary Fig. 4: Marker expression in wild-type and KRAS mutant H6C7 cell line samples.
Supplementary Fig. 5: Comparison of samples cultured in non-adherent versus adherent conditions.
Supplementary Fig. 6: Comparison of PANC-1, CFPAC1 and COLO 357/FG cell line samples cultured in non-adherent versus adherent conditions.
Supplementary Fig. 7: Correlation analysis.
Supplementary Fig. 8: Mass cytometry analysis of patient tissue samples.
Supplementary Fig. 9: Manual gating of marker$^{hi}$ cells in patient tissue samples.
Supplementary Fig. 10: Analysis of patient scRNA-seq data.

#### Supplementary Tables

Supplementary Table 1: Mass cytometry panel.
Supplementary Table 2: Mass cytometry antibody validation.
Supplementary Table 3: Mutational profiles of pancreatic cancer cell lines.
Supplementary Table 4: Clinical information and mutational profiles of patient tissue samples.
Supplementary Table 5: Event counts of mass cytometry samples.

### Supplementary Figures

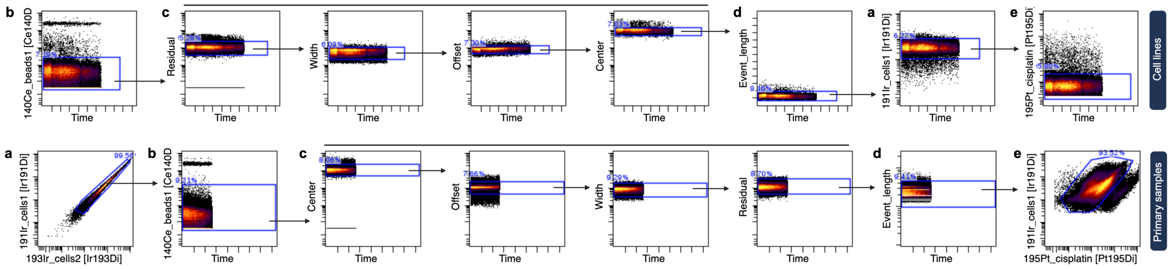


**Supplementary Figure 1: Gating strategy for mass cytometry data clean-up. a** Inclusion of iridium/DNA-positive events. **b** Removal of beads. **c** Gating using Gaussian parameters. **d** Removal of doublets. **e** Removal of cisplatin-positive (i.e., dead) cells.

**a**

**b**

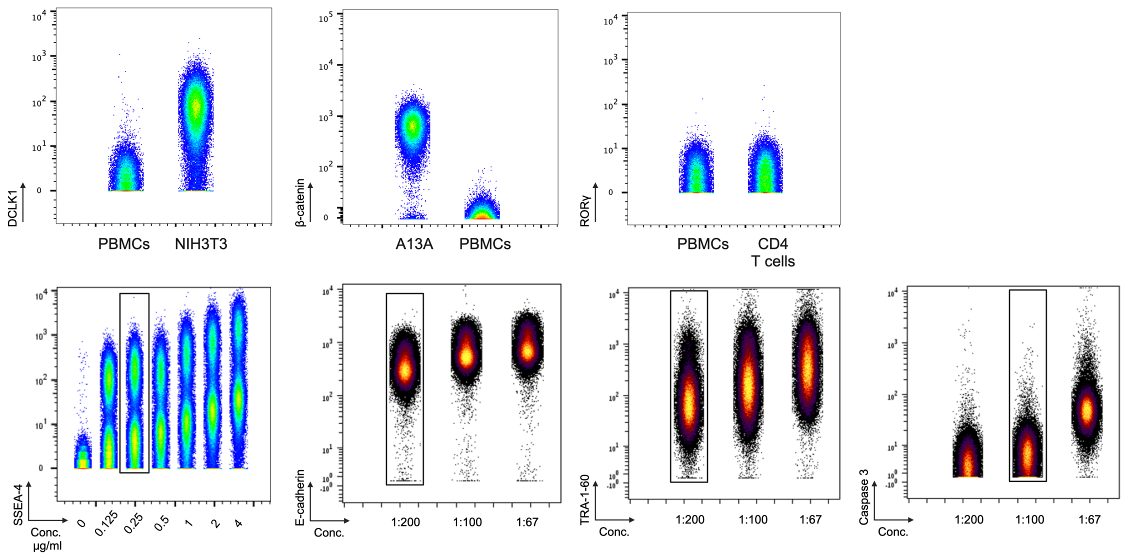


**Supplementary Figure 2: Mass cytometry antibody validation and titration. a** Validation of DCLK1, β-catenin and RORγ antibodies using PBMCs (negative for DCLK1 and β-catenin), pancreatic cancer cell line A13A (positive for β-catenin), mouse embryonic fibroblast line NIH3T3 (positive for DCLK1) and CD4 T cells (contains RORγ-positive subset). **b** Examples of mass cytometry antibody titration. Black rectangle outline indicates the optimal staining concentration. Conc., concentration.

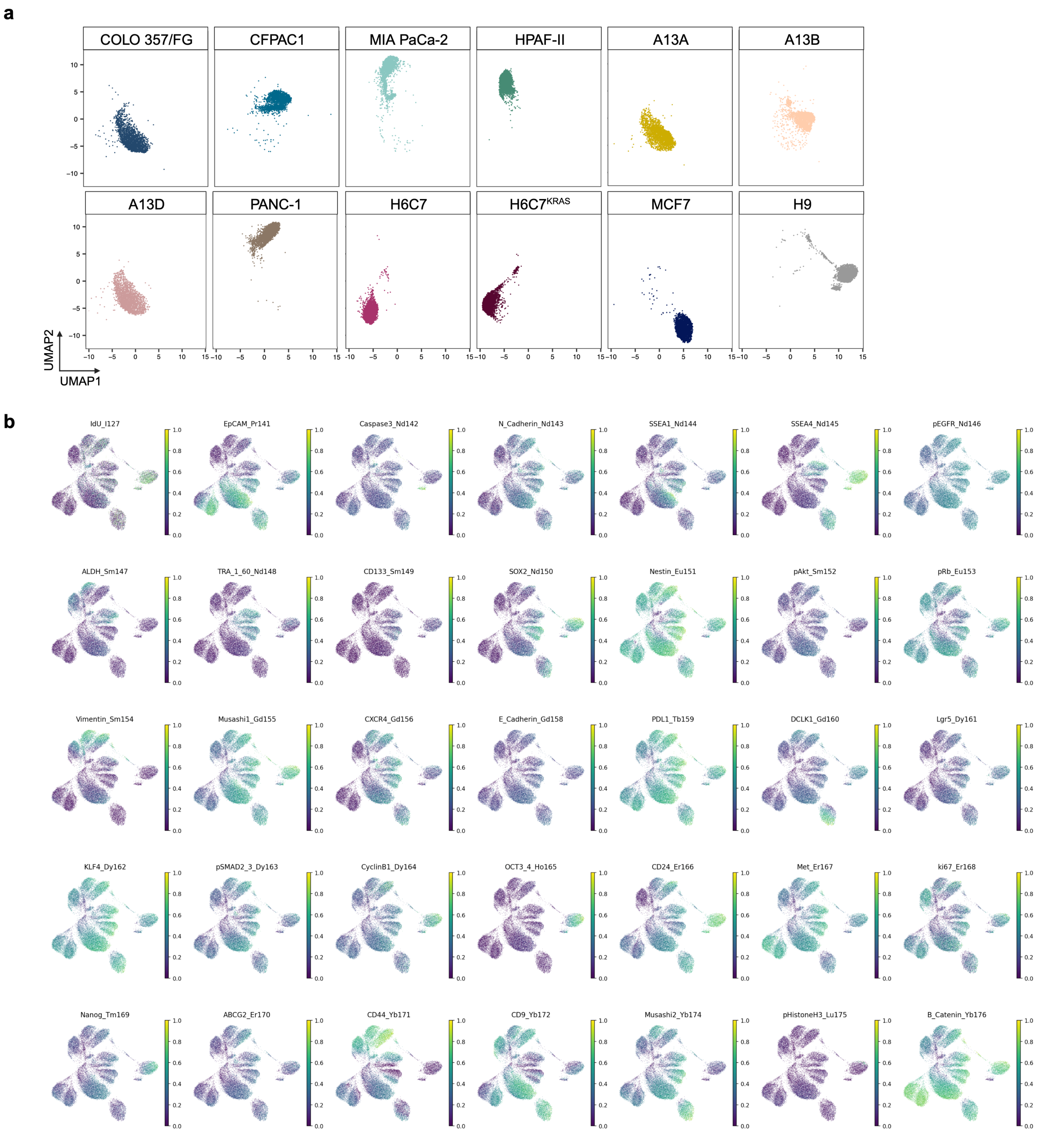


**Supplementary Figure 3: Characterisation of PDAC cell lines cultured in the non-adherent conditions.** **a** UMAP visualisation as in Fig. 2c split by sample. **b** UMAP visualisation as in Fig. 2c coloured by normalised expression of indicated markers. Live, single cells are shown.

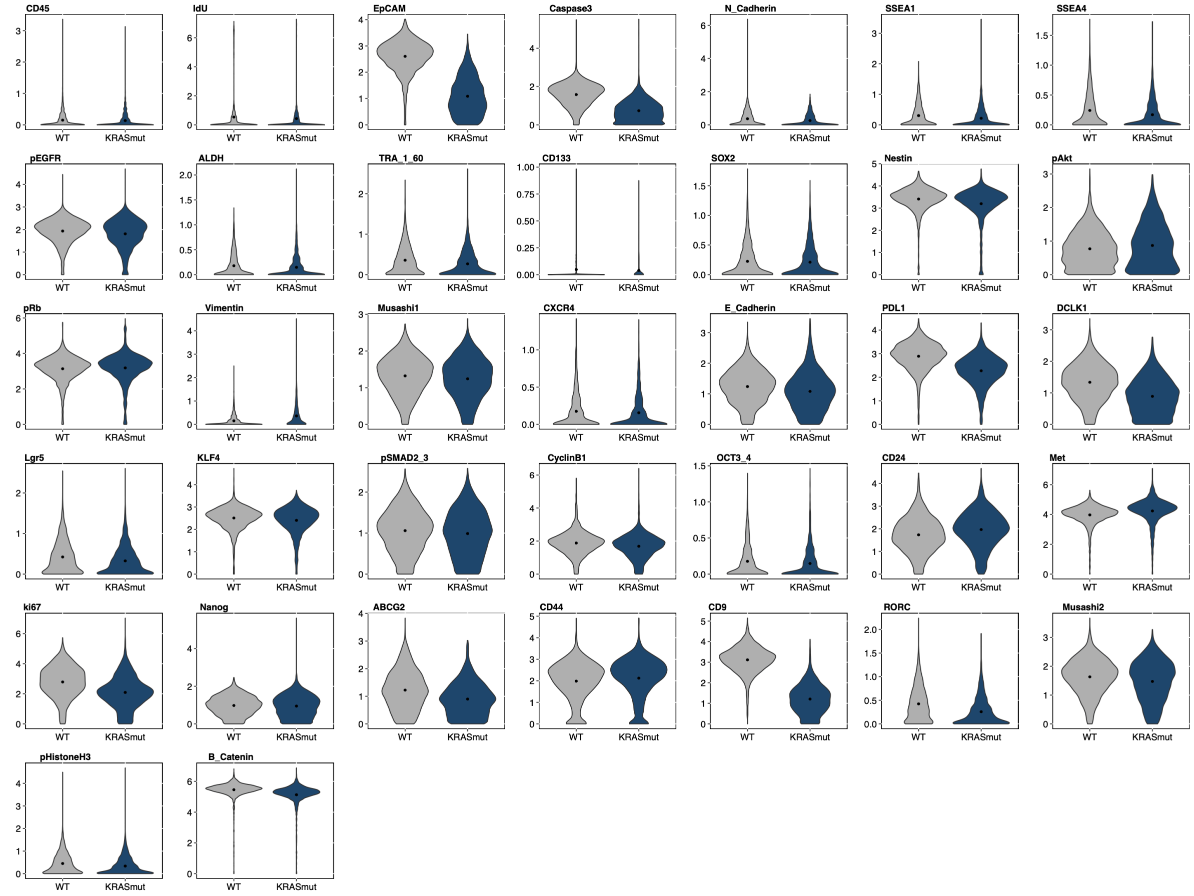


**Supplementary Figure 4: Marker expression in wild-type and KRAS mutant H6C7 cell line samples.** Violin plots showing the indicated marker expression in the samples of wild-type (WT) or KRAS mutant ($KRAS^{mut}$) H6C7 cell line. Point represents the mean intensity.

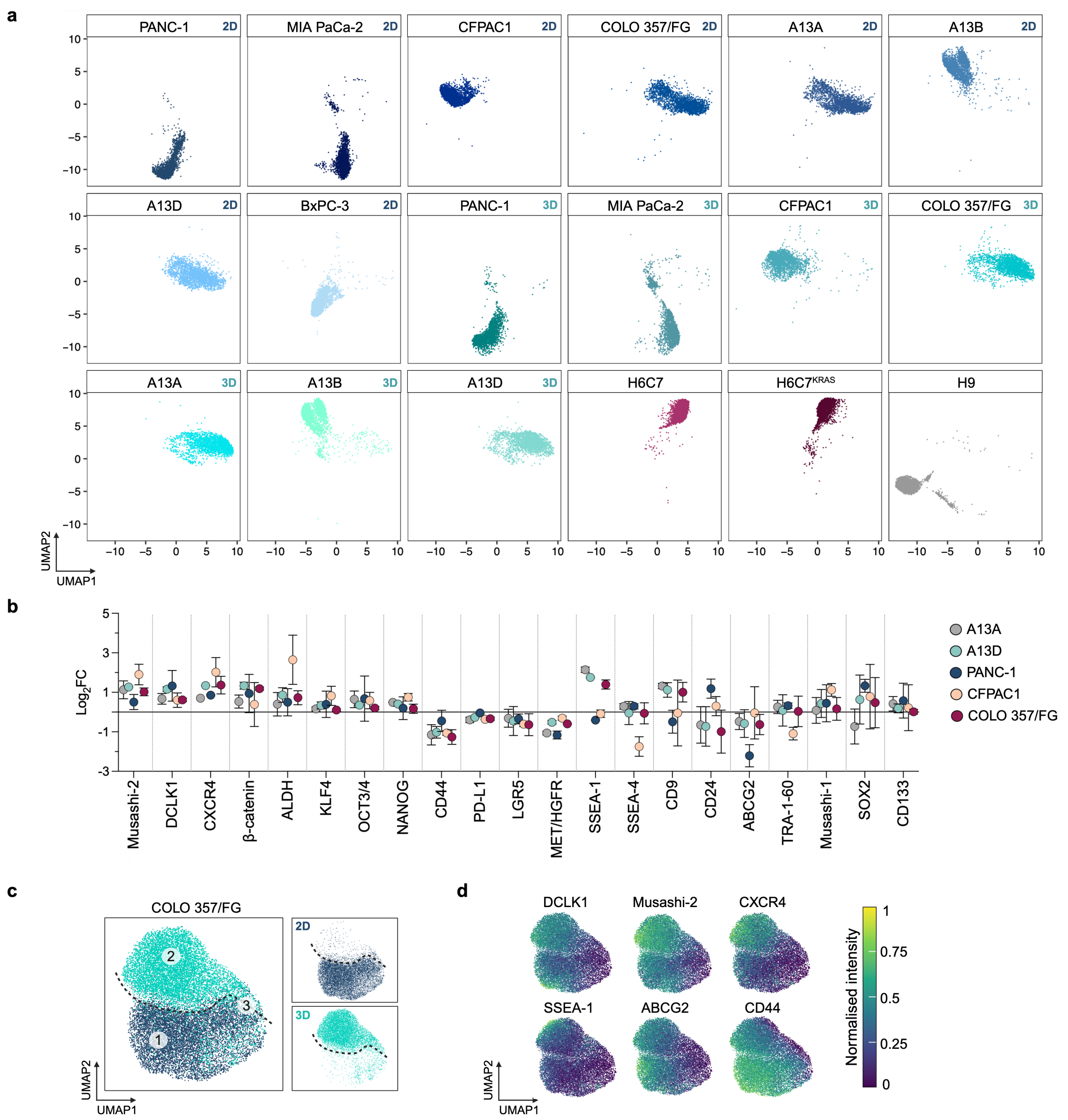


**Supplementary Figure 5: Comparison of samples cultured in non-adherent versus adherent conditions. a** UMAP visualisation as in Fig. 3b split by sample. **b** Relative mean expression levels of indicated markers in non-adherent samples of PDAC cell lines compared to the respective adherent samples. Data are shown as mean ± SD (n = 2). **c** UMAP visualisation of samples of COLO 357/FG cell line from adherent (2D) and non-adherent (3D) culture. Dimensionality reduction was calculated using subsampled data (9,000 cells/sample). Areas 1-3 exhibit distinct proportions of cells from 2D and 3D culture. **d** UMAP visualisation as in **c** coloured by normalised expression of indicated markers**. a, c-d** Live, single cells are shown. FC, fold change.

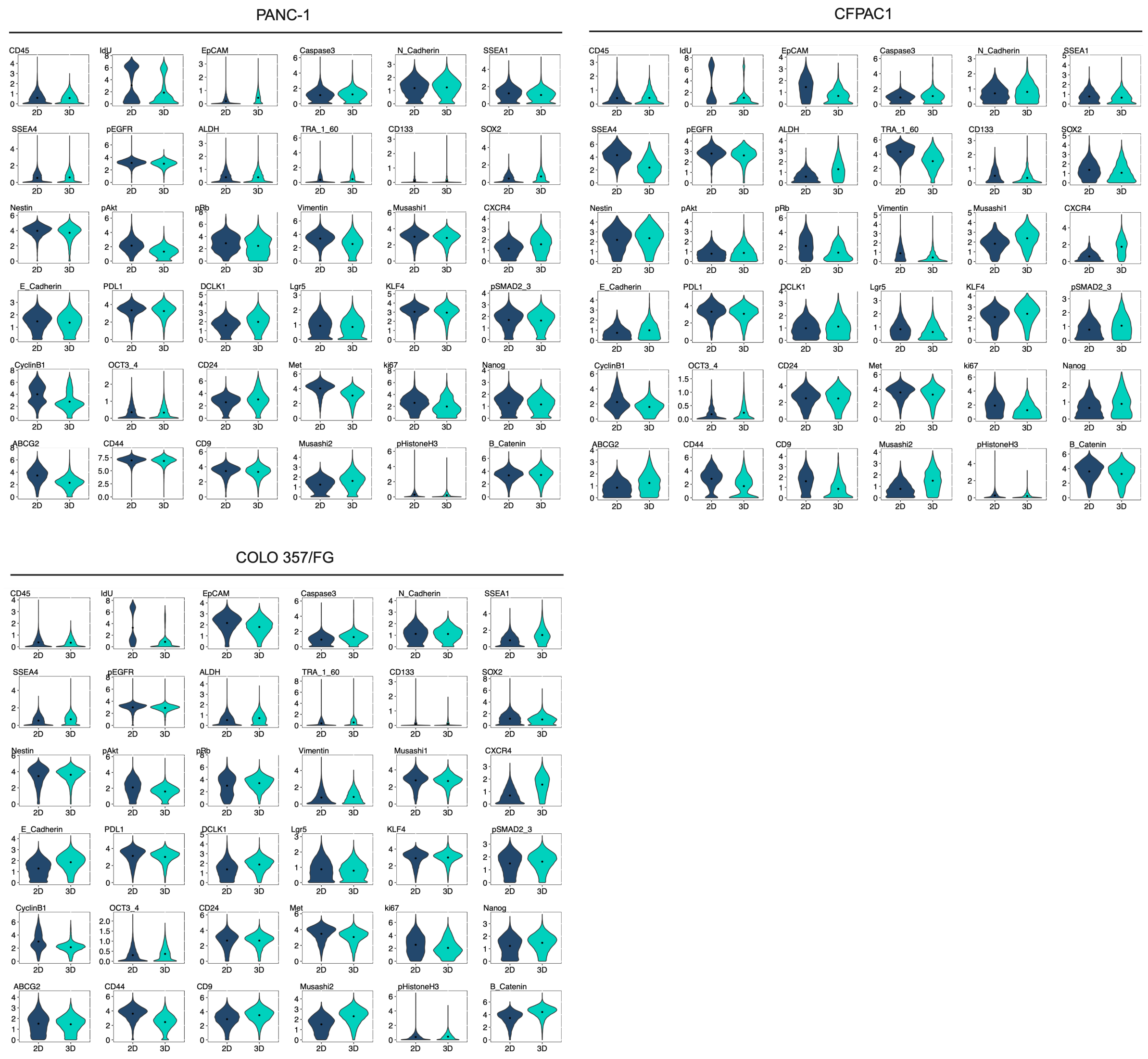


**Supplementary Figure 6: Comparison of PANC-1, CFPAC1 and COLO 357/FG cell line samples cultured in non-adherent versus adherent conditions.** Violin plots showing the indicated marker expression in the samples of PANC-1, CFPAC1 or COLO 357/FG cell line cultured in the adherent (2D) or non-adherent (3D) conditions. Point represents the mean intensity.

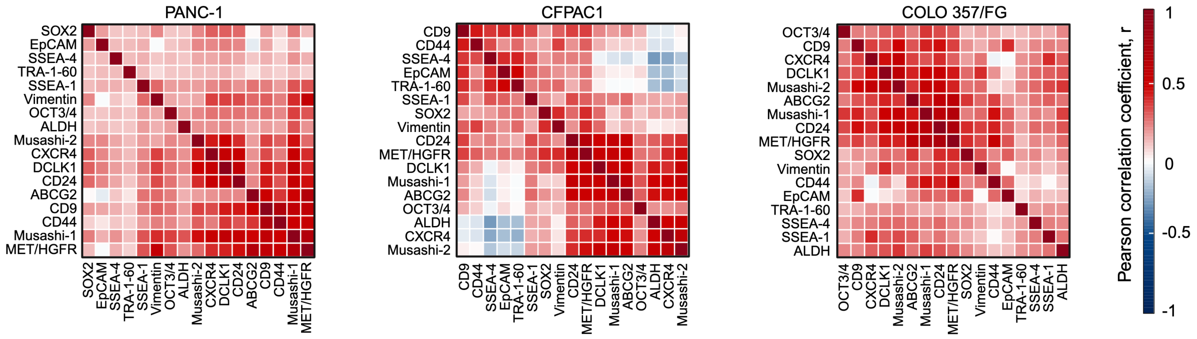


**Supplementary Figure 7: Correlation analysis.** Heatmaps of the correlation coefficients for the indicated markers. Correlation analysis was performed on the non-adherent samples of PANC-1, CFPAC1 and COLO 357/FG cell lines (2,220 cells/sample).

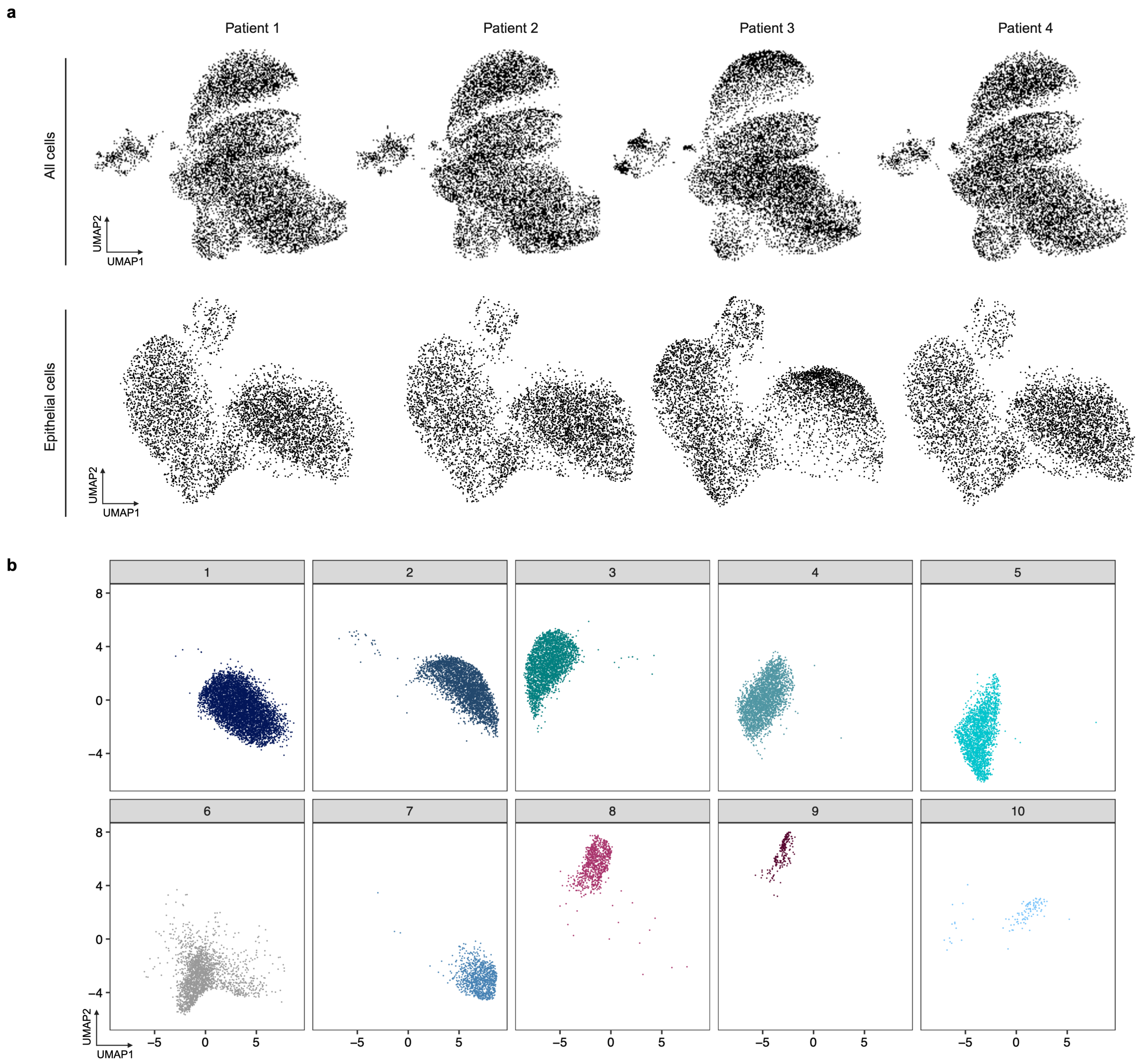


**Supplementary Figure 8: Mass cytometry analysis of patient tissue samples. a** UMAP visualisation as in Fig. 4a (top) and Fig. 4d (bottom) split by patient. **b** UMAP visualisation as in Fig. 4d split by Phenograph cluster. Each dot represents a cell. Live, single cells are shown.

**a**

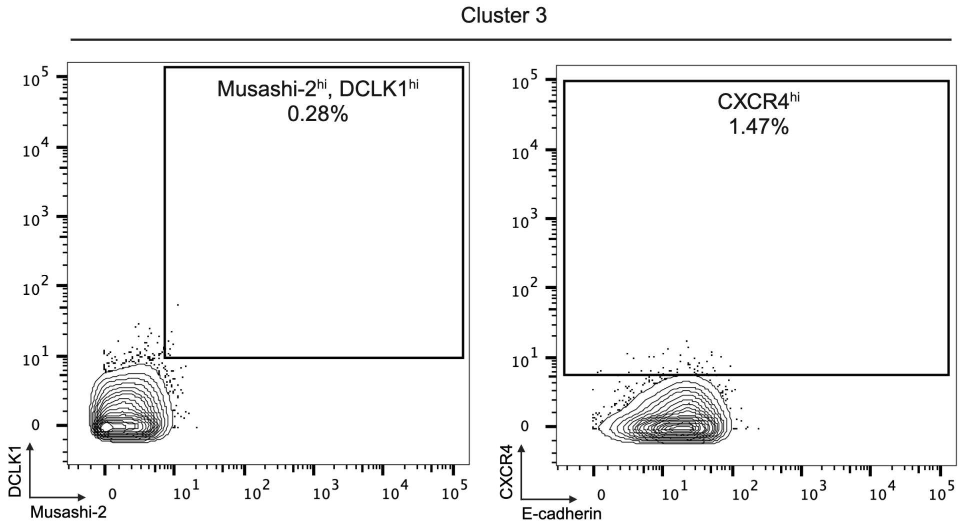


**b**

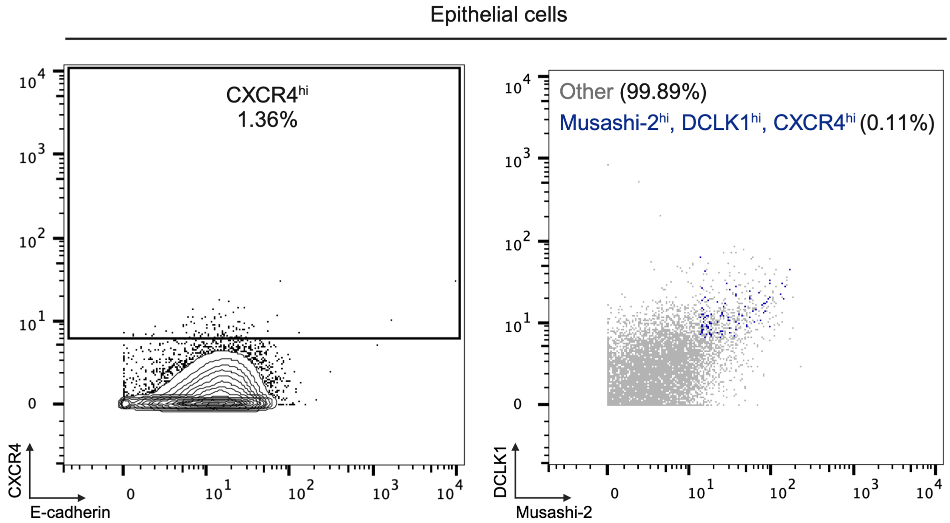


**Supplementary Figure 9: Manual gating of marker$^{hi}$ cells in patient tissue samples. a** Examples of marker$^{hi}$ gates in the Phenograph cluster 3. **b** Two-dimensional plots illustrating low abundance of CXCR4$^{hi}$ and triple-high (Musashi-2$^{hi}$, DCLK1$^{hi}$ and CXCR4$^{hi}$) epithelial cells in the patient tissue samples. Live, single cells are shown.

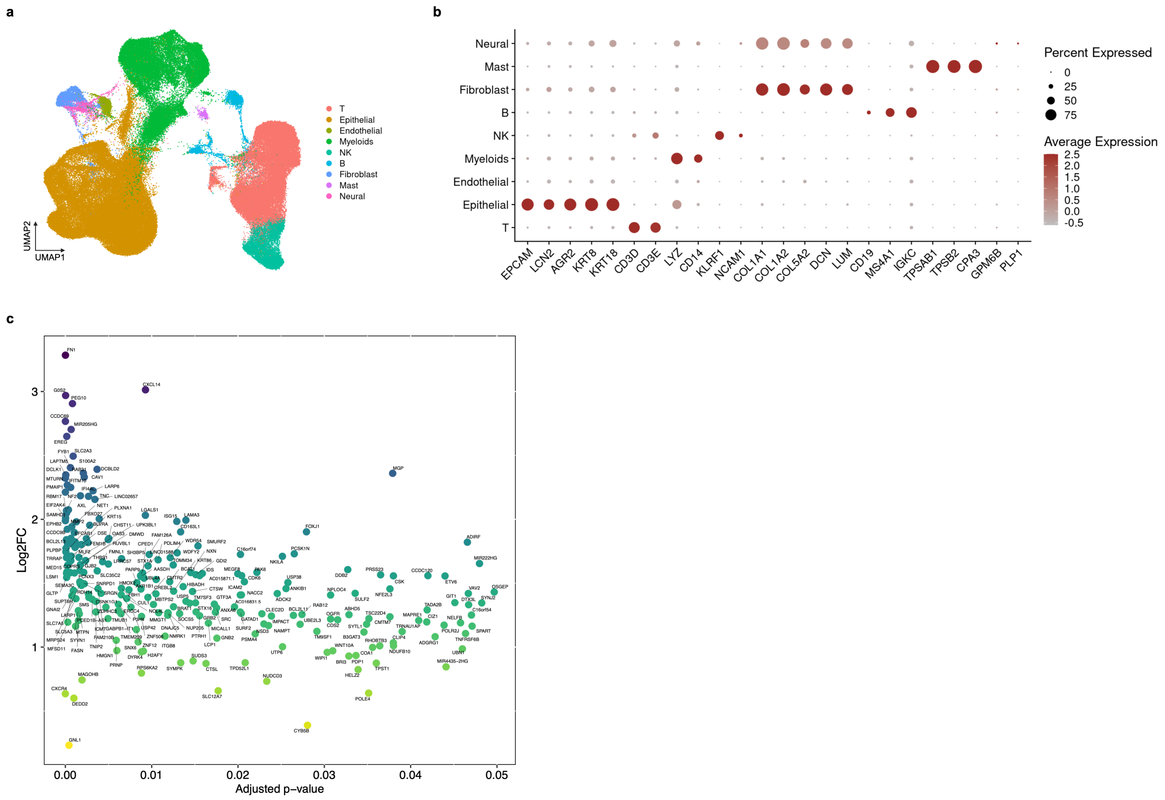


**Supplementary Figure 10: Analysis of patient scRNA-seq data. a** UMAP visualisation of scRNA-seq data. Dimensionality reduction was calculated on highly variable features. Each dot represents a cell and is coloured by the cell type. **b** Expression of marker genes by annotated cell type clusters. **c** Genes significantly (adjusted p-value <0.05) upregulated in triple-high (Musashi-2$^{hi}$, DCLK1$^{hi}$ and CXCR4$^{hi}$) cells compared to the rest of epithelial cells. FC, fold change.

**Supplementary Table 1: Mass cytometry panel.**

| Target | Channel | Company | Catalogue | Clone | Label | Conc. |
|---|---|---|---|---|---|---|
| Barcode | 102Pd<br>104Pd<br>105Pd<br>106Pd<br>108Pd<br>110Pd | Standard BioTools | 201060 | - | - | - |
| CD45 | 89Y | Standard BioTools | 3089003B | HI30 | Commercial | 1:200 |
| IdU | 127I | Standard BioTools | 201127 | - | - | - |
| EpCAM/CD326/ESA | 141Pr | Standard BioTools | 3141006B | 9C4 | Commercial | 1:1000 |
| Cleaved caspase 3 | 142Nd | Standard BioTools | 3142004A | D3E9 | Commercial | 1:100 |
| N-cadherin | 143Nd | Standard BioTools | 3143016B | 8C11 | Commercial | 1:100 |
| SSEA-1/CD15 | 144Nd | Standard BioTools | 3144019B | W6D3 | Commercial | 1:100 |
| SSEA-4 | 145Nd | BD Biosciences | 560073 | MC813-70 | In-house | 0.5 µg/ml |
| Phospho-EGFR (Y1068) | 146Nd | Standard BioTools | 3146007A | D7A5 | Commercial | 1:100 |
| ALDH | 147Sm | Standard BioTools | 3147015B | 44/ALDH | Commercial | 1:100 |
| TRA-1-60 | 148Nd | Standard BioTools | 3148012B | TRA160 | Commercial | 1:200 |
| CD133 | 149Sm | BioLegend | 397902 | W6B3C1 | In-house | 0.125 µg/ml |
| SOX2 | 150Nd | Standard BioTools | 3150019B | O30678 | Commercial | 1:100 |
| Nestin | 151Eu | Standard BioTools | 3151013A | 25/Nestin | Commercial | 1:100 |
| Phospho-AKT (S473) | 152Sm | Standard BioTools | 3152005A | D9E | Commercial | 1:200 |
| pRb | 153Eu | BD Biosciences | 558389 | J112-906 | In-house | 0.25 µg/ml |
| Vimentin | 154Sm | Standard BioTools | 3154014A | D21H3 | Commercial | 1:200 |
| Musashi-1 | 155Gd | Standard BioTools | 3155013B | 14H1 | Commercial | 1:100 |
| CXCR4 | 156Gd | Standard BioTools | 3156029B | 12G5 | Commercial | 1:200 |
| E-cadherin | 158Gd | Standard BioTools | 3158021A | 24E10 | Commercial | 1:400 |
| PD-L1 | 159Tb | Standard BioTools | 3159029B | 29E.2A3 | Commercial | 1:100 |
| DCLK1 | 160Gd | Abnova | H00009201-M03 | 6H4 | In-house | 0.5 µg/ml |
| LGR5 | 161Dy | Standard BioTools | 3161025B | 4D11F8 | Commercial | 1:100 |
| KLF4 | 162Dy | Standard BioTools | 3162022A | D1F2 | Commercial | 1:100 |

| | | | | | | |
|---|---|---|---|---|---|---|
| Phospho-SMAD2/3 | 163Dy | Cell Signaling | 8828 | D27F4 | In-house | 0.25 µg/ml |
| Cyclin B1 | 164Dy | Standard BioTools | 3164010A | GNS1 | Commercial | 1:100 |
| OCT3/4 | 165Ho | Standard BioTools | 3165023A | 40/Oct3 | Commercial | 1:100 |
| CD24 | 166Er | Standard BioTools | 3166007B | ML5 | Commercial | 1:100 |
| MET/HGFR | 167Er | Standard BioTools | 3167017A | D1C2 | Commercial | 1:200 |
| Ki-67 | 168Er | Standard BioTools | 3168007B | B56 | Commercial | 1:100 |
| NANOG | 169Tm | Standard BioTools | 3169014A | N31-355 | Commercial | 1:100 |
| ABCG2 | 170Er | R&D Systems | MAB995 | 5D3 | In-house | 1 µg/ml |
| CD44 | 171Yb | Standard BioTools | 3171003B | IM7 | Commercial | 1:100 |
| CD9 | 172Yb | Standard BioTools | 3172010B | SN4 C33A2 | Commercial | 1:200 |
| RORγ | 173Yb | R&D Systems | MAB6109 | 600214 | In-house | 1 µg/ml |
| Musashi-2 | 174Yb | R&D Systems | MAB3255 | 960121 | In-house | 1 µg/ml |
| Phospho-histone H3 (S28) | 175Lu | Standard BioTools | 3175012A | HTA28 | Commercial | 1:1000 |
| β-catenin | 176Yb | Abcam | ab242424 | SP328 | In-house | 0.5 µg/ml |
| Iridium (cells) | 191/193Ir | Standard BioTools | 201192A | - | - | - |
| Cisplatin (live/dead) | 195Pt | Standard BioTools | 201064 | - | - | - |

**Supplementary Table 2: Mass cytometry antibody validation.**

| Target | Negative/low cells | Positive/high cells | Status |
|---|---|---|---|
| SSEA-4 | A13D | H9 | Validated |
| CD133 | A13D | H9 | Validated |
| pRb | PBMCs | A13A | Validated |
| DCLK1 | PBMCs | NIH3T3 | Validated |
| Phospho-SMAD2/3 | A13A (SB431542-treated) | A13A (activin A-treated) | Poor performance |
| ABCG2 | A13A | PANC-1 | Validated |
| RORγ | PBMCs | CD4 T cells | Not validated |
| Musashi-2 | - | HeLa and HEK293T | Validated |
| β-catenin | A13A and PBMCs | A13A | Validated |

**Supplementary Table 3: Mutational profiles of pancreatic cancer cell lines.**

| Cell line | *KRAS* | *TP53* | *CDKN2A* | *SMAD4* | Reference |
|---|---|---|---|---|---|
| A13A | G12V | WT | ND | WT | (Embuscado et al., 2005) |
| A13B | G12V | WT | ND | WT | (Embuscado et al., 2005) |
| A13D | G12V | WT | ND | WT | (Embuscado et al., 2005) |
| COLO 357/FG | G12D | WT | WT | WT | (Kleeff & Korc, 1998; Rückert F, 2013) |
| PANC-1 | G12D | R273H | HD | WT | (Deer et al., 2010) |
| MIA PaCa-2 | G12C | R248W | HD | WT | (Deer et al., 2010) |
| HPAF-II | G12D | P151S | DEL | WT | (Deer et al., 2010) |
| CFPAC1 | G12V | Mut. | Methylated | Mut. | (Deer et al., 2010) |
| BxPC-3 | WT | Y220C | HD | HD | (Deer et al., 2010) |

DEL, deletion; HD, homozygous deletion; Mut., mutation; ND, not determined; WT, wild-type.

**Supplementary Table 4: Clinical information and mutational profiles of patient tissue samples.**

| Patient | Diagnosis | Sex | Age | TNM Stage | Treatment history | Mutational profile | | | |
|---|---|---|---|---|---|---|---|---|---|
| | | | | | | *KRAS* | *TP53* | *CDKN2A* | *SMAD4* |
| Patient 1 | PDAC | Female | 73 | T2N1 (L1 V1 R0) | Neoadjuvant chemotherapy | G12D | DEL | M52K | V354L |
| Patient 2 | PDAC | Male | 63 | T2N1Mx | Neoadjuvant chemotherapy | G12V | R273H | WT | V354L |
| Patient 3 | PDAC | Male | 76 | T3N1M0 | 6 cycles of Folfirinox | G12D | DEL | M52K | WT |
| Patient 4 | PDAC | Female | 59 | T2N2Mx | 5 cycles of Folfirinox | G12D | R248W | WT | WT |

DEL, deletion; WT, wild-type.

**Supplementary Table 5: Event counts of mass cytometry samples.**

**Cell line samples** (live, single cells)

| Sample | Replicate 1 | Replicate 2 |
|---|---|---|
| A13A_adh_1 | 123263 | 88826 |
| A13A_adh_2 | 200711 | 60778 |
| A13A_non_adh | 223709 | 49026 |
| A13B_adh | 110866 | 70599 |
| A13B_non_adh | 2926 | 29364 |
| A13D_adh | 168743 | 85448 |
| A13D_non_adh | 68779 | 113633 |
| PANC-1_adh | 57043 | 76869 |
| PANC-1_non_adh | 11098 | 62281 |
| CFPAC1_adh | 266278 | 87236 |
| CFPAC1_non_adh | 74323 | 9307 |
| MIA_PaCa-2_adh | 189789 | 113849 |
| MIA_PaCa-2_non_adh | 66650 | 3850 |
| COLO357/FG_adh | 139494 | 144459 |
| COLO357/FG_non_adh | 95853 | 72193 |
| HPAF-II_adh | 179307 | 87957 |
| HPAF-II_non_adh | 293316 | 57941 |
| BxPC-3 | 275798 | 85858 |
| MCF7_adh | 213984 | 60795 |
| MCF7_non_adh | 292891 | 118706 |
| H6C7 | 137102 | 4288 |
| $H6C7^{KRAS}$ | 3241 | 2272 |
| H9 | 96688 | 86123 |

**Patient tissue samples**

| Sample | Live, single cells |
|---|---|
| Patient 1 | 200685 |
| Patient 2 | 135999 |
| Patient 3 | 38803 |
| Patient 4 | 120328 |
